# AGAL misprocessing-induced ER stress and the unfolded protein response: lysosomal storage-independent mechanism of Fabry disease pathogenesis?

**DOI:** 10.1101/2022.09.27.509714

**Authors:** Martina Živná, Gabriela Dostálová, Veronika Barešová, Dita Mušálková, Ladislav Kuchař, Befekadu Asfaw, Helena Poupětová, Hana Vlášková, Tereza Kmochová, Petr Vyletal, Hana Hartmannová, Kateřina Hodaňová, Viktor Stránecký, Lenka Steiner-Mrázová, Aleš Hnízda, Martin Radina, Miroslav Votruba, Jana Sovová, Helena Trešlová, Larisa Stolnaja, Petra Reková, Lenka Roblová, Eva Honsová, Helena Hůlková, Ivan Rychlík, Anthony J. Bleyer, Aleš Linhart, Jakub Sikora, Stanislav Kmoch

**Affiliations:** Research Unit for Rare Diseases, Department of Paediatrics and Inherited Metabolic Disorders, First Faculty of Medicine, Charles University in Prague and General University Hospital in Prague, Czech Republic; Second Department of Internal Cardiovascular Medicine, First Faculty of Medicine, Charles University and General University Hospital, Prague, Czech Republic; Diagnostic laboratory, Department of Pediatrics and Inherited Metabolic Disorders, General University Hospital, Prague, Czech Republic; Institute of Pathology, First Faculty of Medicine, Charles University and General University Hospital, Prague, Czech Republic; Department of Neurology and Centre of Clinical Neuroscience, First Faculty of Medicine, Charles University and General University Hospital in Prague, Czech Republic; AeskuLab Pathology, Prague, Czech Republic; Department of Medicine, Third Faculty of Medicine, Charles University in Prague and Faculty Hospital Kralovske Vinohrady, Prague, Czech Republic; Section on Nephrology, Wake Forest School of Medicine, Medical Center Blvd., Winston-Salem, NC, USA

**Keywords:** Fabry disease, pathogenesis, ER stress, unfolded protein respose

## Abstract

**Background:** Classic Fabry disease (FD) is caused by *GLA* mutations that result in enzymatic deficiency of alpha-galactosidase A (AGAL), lysosomal storage of globotriaosylceramide, and a resulting multisystemic disease. In non-classic later-onset FD, patients have some preserved AGAL activity and a milder disease course, though female carriers may also be affected. While FD pathogenesis has been mostly attributed to catalytic deficiency of mutated AGAL, lysosomal storage and impairment of lysosomal functions, other pathogenic factors may be important, especially in non-classic later-onset FD.

**Methods:** We characterized the clinical, biochemical, genetic, molecular, cellular and organ pathology correlates of the p.L394P AGAL variant that was identified in six individuals with end-stage kidney disease by the Czech national screening program for FD and by further screening of 25 family members.

**Results:** Clinical findings revealed a milder clinical course with ~15% residual AGAL activity. Laboratory investigations documented intracellular retention of mutated AGAL with resulting ER stress and the unfolded protein response (UPR). Kidney biopsies did not show lysosomal storage. We observed similar findings of ER stress and UPR with several other classic and non-classic FD missense *GLA* variants.

**Conclusions:** We identified defective proteostasis of mutated AGAL resulting in chronic ER stress and UPR of AGAL expressing cells (hereafter referred to as AGALopathy) as an important contributor to FD pathogenesis. These findings provide insight into non-classic later-onset FD and may better explain clinical manifestations with implications for pathogenesis, clinical characterization and treatment of all FD forms.

**Significance statement:** Catalytic deficiency of mutated AGAL is responsible for classicFabry disease (FD) pathogenesis but does not fully explain the findings in non-classic later-onset FD, in which affected individuals and female carriers develop clinical manifestations despite some AGAL activity and variably mitigated lysosomal storage. In this investigation of individuals with the p.L394P AGAL variant, we identified defective proteostasis of mutated AGAL resulting in chronic endoplasmic reticulum stress and the unfolded protein response as significant contributors to pathogenesis of non-classic later-onset FD. Similar effects were documented also in other AGAL variants identified in classic and non-classicFD. Endoplasmic reticulum stress and the unfolded protein response therefore play an important role in FD.

## Introduction

Fabry disease (FD; OMIM #301500) is an X-linked condition caused by pathogenic variants of the *GLA* gene that result in the absence or enzymatic deficiency of alpha-galactosidase A (AGAL). This enzyme defect leads to lysosomal storage of globotriaosylceramide (Gb3Cer) in a variety of cell types throughout the body and manifests as a multisystemic disease affecting the heart, kidney, blood vessels, peripheral nervous system, skin, and eyes.

Classic FD, is best exemplified by males with truncating or missense *GLA* variants who have no AGAL enzyme activity, increased Gb3Cer concentrations in plasma and characteristic lysosomal storage vacuoles in affected cell types/tissues. These individuals typically develop cardiac, kidney and neurologic complications in the 3^rd^ or 4^th^ decade of life [1]. Many, but not all, patients respond to enzyme replacement therapy with recombinant AGAL [2], though the response is sub-optimal [3]. Migalastat, a small-molecule chaperone that facilitates AGAL trafficking from the endoplasmic reticulum to lysosomes for certain (amenable) nontruncated enzymes, is also helpful in some cases [4].

Over the last two decades an increasing number of patients have been described with non-classic later-onset FD [5]. Many of these individuals have been identified by genetic screening of newborns or CKD populations [6], allowing the identification of new *GLA* variants and corresponding clinical manifestations [7, 8]. These individuals typically have missense *GLA* variants, varying levels of AGAL activity and a modest increase in plasma Gb3Cer. The phenotype is clinically more homogeneous with disease occurring later in life and often limited to cardiac or kidney disease [9]. Lysosomal storage is less constant and may be less pronounced than in classic FD. Manifestations of FD have also been reported in females heterozygous for pathogenic *GLA* variants. Clinical symptoms occur with later age at onset than in males and often do not correlate with plasma AGAL activity [10], with higher rates of symptomatic disease than expected by random X-inactivation [11].

While the pathogenesis of FD has been mostly attributed to catalytic deficiency of mutated AGAL, lysosomal Gb3Cer accumulation and impairment of lysosomal functions [12], other factors may also contribute [13–15].

For example, endoplasmic reticulum (ER) misfolding and intracellular retention of mutant proteins resulting in ER stress and the unfolded protein response (UPR) play a central pathogenic role in many human diseases [16].

AGAL is synthesized in the ER, and post-translationally modified and delivered to the lysosome through the secretory pathway [17]. Specific *GLA* variants might affect the synthesis, processing, and stability of AGAL in different ways [18]. While misfolding of some AGAL variants and their accelerated degradation through endoplasmic reticulum associated protein degradation (ERAD) constitute the established mechanism leading to lysosomal enzyme deficiency [19], the potential contribution of AGAL missprocessing, intracellular retention and consequent ER stress and UPR to development of FD symptoms has not yet been investigated.

In a recently completed nationwide FD screening program in patients receiving maintenance dialysis therapy, we identified six probands with the *GLA* c.1181T>C (p.L394P AGAL) variant. Family screening led to the identification of total of 31 carriers of the same variant. This family cohort had a high prevalence of progressive kidney disease more related to cell and tissue damage caused by intracellular retention of the mutated AGAL than to the actual enzymatic deficiency and resulting excess lipid storage in the kidney. Laboratory investigations revealed intracellular retention of mutated AGAL with resulting ER stress and the UPR. We then characterized and observed similar manifestation with several other missense *GLA* variants identified in individuals with classic and non-confirmed the expectedclassic FD.

These clinical and laboratory studies identify defective proteostasis of mutated AGAL resulting in chronic ER stress and UPR of AGAL expressing cells (hereafter referred to as AGALopathy) as important etiologic and lysosomal storage-independent factors with implications for pathogenesis, clinical characterization and treatment of all FD forms.

## Results

### Patients

As part of the Czech nationwide FD screening program of individuals with end-stage kidney disease, the six probands were identified as having ~15% residual blood AGAL activity and the c.1181T>C *GLA* variant, encoding missense p.L394P AGAL. Family histories revealed numerous individuals with chronic kidney disease (**Figure 1**). Whole genome sequencing performed in F1_III.5 and F3_III.1 revealed that these individuals shared 4.7 Mb of genetic material on chromosome X (chrX: 100016210 – 104743750; (hg19)), confirmed the expected founder effect and excluded other potential disease-causing variants that either might be linked to the c. 1181T>C variant or localized elsewhere in the genome. The variant had not been previously reported in population genetic databases, is conserved across species, and is classified as pathogenic by various pathogenicity classifiers.

**Figure 1.**
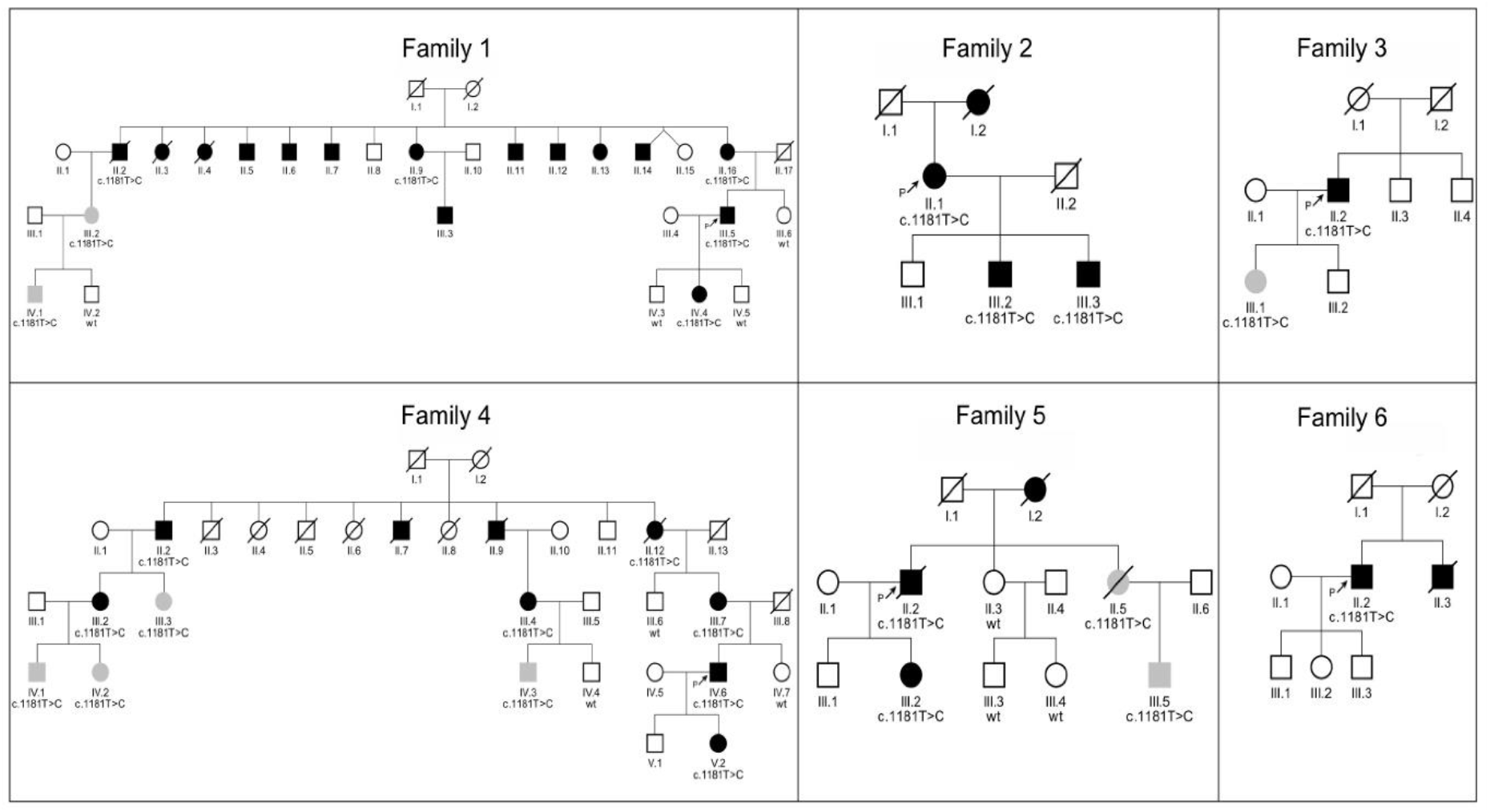
Family pedigrees. Generations and individuals are identified successively by Roman and Arabic numerals, respectively. Black symbols denote individuals with at least one pre-specified FD manifestation. Gray symbols denote genetically affected individuals with no FD manifestation. Open symbols denote clinically and genetically unaffected individuals. The c. 1181T>C or wt denote presence or absence of the *GLA* mutation, respectively; if not stated indicates that DNA was unavailable for investigation.

Family screening identified 30 carriers (15 males and 15 females) with the same c.1181T>C variant and a high prevalence of FD symptoms (**Supplemetary Table S1**). Laboratory investigations revealed that all of 7 investigated male carriers had AGAL activities below the normal range in plasma and leukocytes and minimally increased plasma lyso-Gb3Cer levels. All but one of the 9 female carriers had normal leukocyte AGAL activities; with two of them having AGAL activities below the normal range in plasma. All female carriers had normal plasma lyso-Gb3Cer levels.

Twenty three (75 %) carriers had at least one pre-specified FD manifestation. The most frequent findings included the presence of proteinuria (n=17), decreased estimated glomerular filtration rate (n=10, including 7 patients with end-stage kidney disease), gastrointestinal symptoms (n=10), ocular changes (n=7), left ventricular hypertrophy LVH (n=7), polyneuropathy (n=8), tremor (n=6) and stroke/TIA history (n=3). Only one carrier (a female) had typical Fabry-related angiokeratomas. Hypertension was present in 9 individuals. Of interest, 12 affected individuals had diabetes as compared to the absence of diabetes in unaffected family members. Biochemical cell-loading studies revealed that the degradation of C23:0,d18:1 globotriaosylceramide (C23Gb3Cer) was reduced in cultured skin fibroblasts of the male with the p.L394P AGAL, but to a lesser extent than in fibroblasts of the male with the classic FD (**Figure 2**). Histopathologic and immunohistochemical studies in kidney biopsies of males with the p.L394P AGAL revealed enlarged glomeruli without proliferation and mild focal interstitial fibrosis. Characteristic lysosomal storage abnormalities found in FD (**Supplementary Figure S1**) were not detected in glomeruli. Ultrastructural electron microscopic studies in kidney biopsies of males with the p.L394P AGAL showed partial effacement of podocyte pedicels and non-membrane bound lipid droplets in proximal tubular epithelium and interstitial cells (some of them likely macrophages). Lysosomal storage typical for FD was not found in any cell type present in the tested tissue at the ultrastructural level (**Figure 3**).

**Figure 2.**
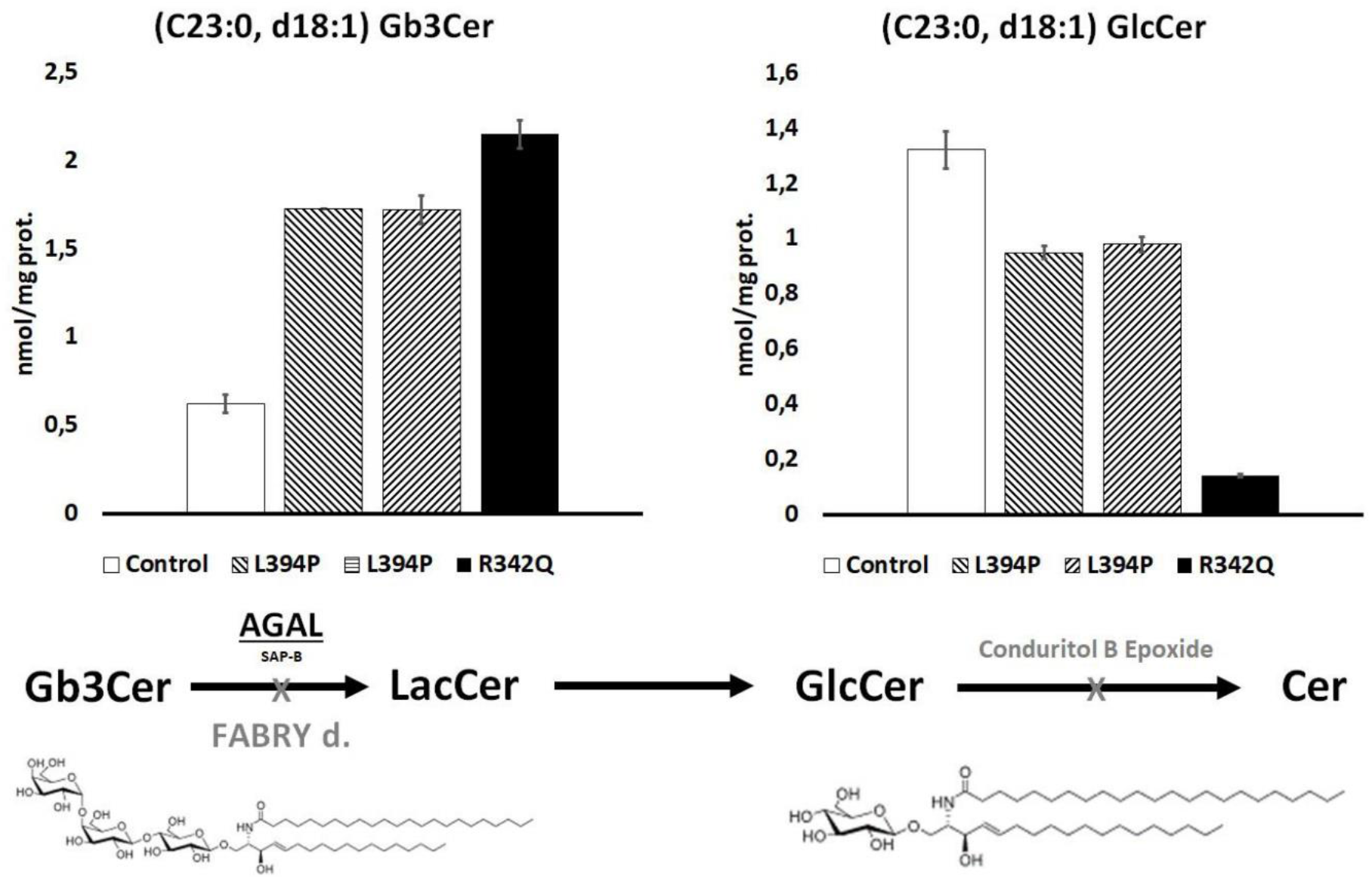
Degradation of globotriaosylceramide in cultured skin fibroblasts. Skin fibroblasts were loaded with mass-labeled C23:0,d18:1 globotriaosylceramide (C23Gb3Cer). Conduritol B epoxide, a covalent inhibitor of GIcCer-ß-Glucosidase, was applied to block the downstream metabolic conversion of glucosylceramide to ceramide. After 4 days of incubation, the original substrate, C23Gb3Cer, and the resulting product - C23:0,d18:1 glucosylceramide (C23GIcCer) were isolated from cell homogenates and quantified by FIA-ESI-MS/MS. Compare to control cells, degradation of the C23Gb3Cer was reduced in cells from two males with the p. L394P AGAL, but to a lesser extent than in cells of the male with classic FD and p.R342Q AGAL.

**Figure 3.**
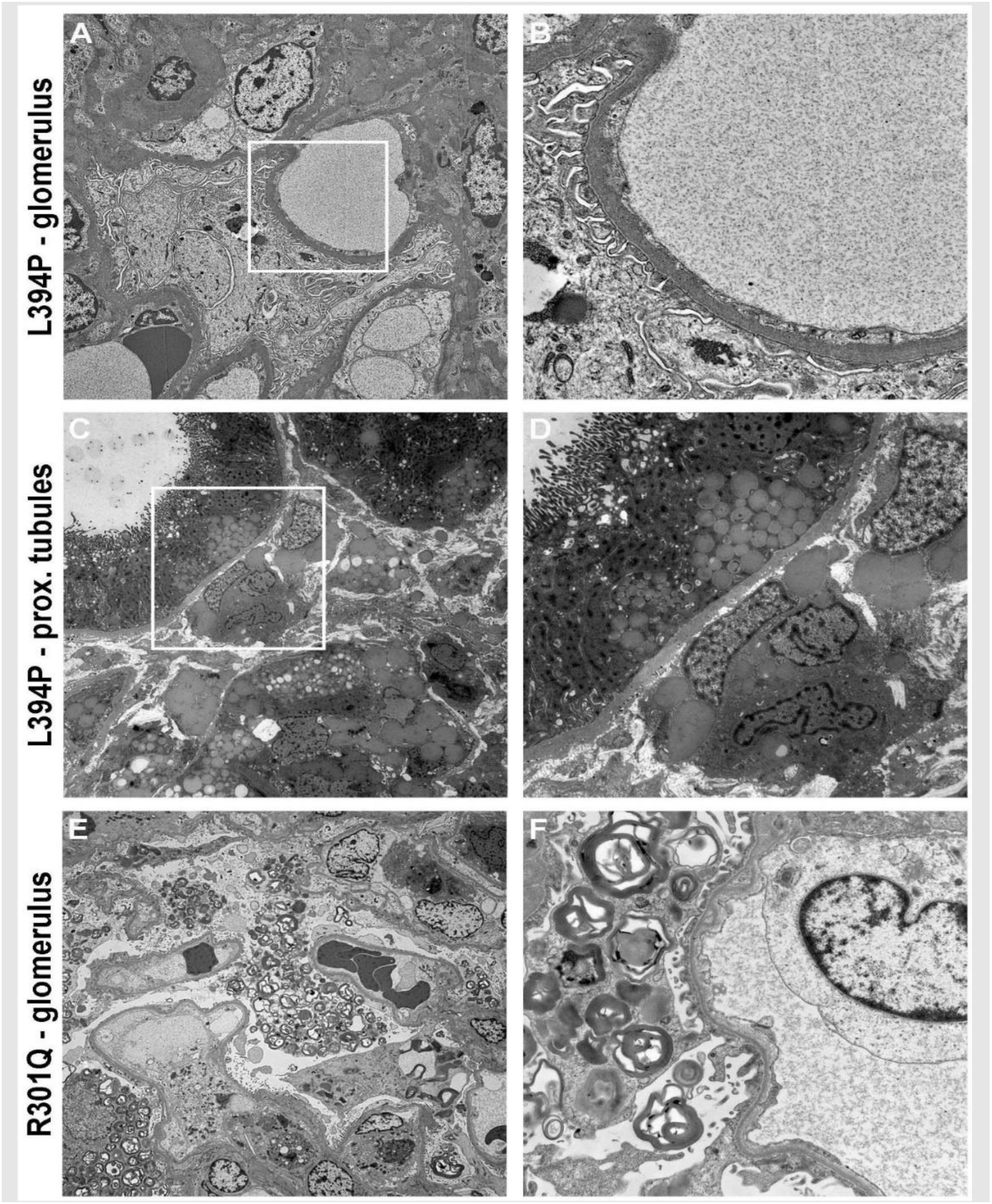
Renal ultrastructural pathology. (A,B) Glomerular structures in a male with the p.L394P AGAL. Podocytes and endothelial cells do not contain typical FD storage lysosomes. Partial pedicel effacement is shown in higher detail in B (black arrows). The white square in A corresponds to the area shown in B. (C,D) Non-mebrane bound lipid droplets (white arrows) in proximal tubular epithelia and interstitial cells (white arrowheads) of a male with the p.L394P AGAL. Notably, lysosomal storage typical for FD was not found in any cell type present in the tested tissue. The white square in C corresponds to the area shown in D. (E,F) Characteristic multilamellar storage lysosomes (black arrows) are abundant in podocytes of a male with classic FD and the p.R3OlQ AGAL. The white square in E corresponds to the area shown in F. Scale bars: A, D, E - 2 mm; C - 5 mm; B, F - 750 nm.

Immunofluorescence studies in kidney biopsies of FD patients and controls showed the usual distribution of AGAL with high expression in tubular epithelial cells and minimal expression in podocytes [20]. However, analyses of the p.L394P AGAL kidney revealed that the mutated AGAL localizes mainly to the ER, endoplasmic reticulum-Golgi intermediate compartment (ERGIC) and co-localizes with transmembrane P24 Trafficking Protein 9 (TMED9), the vesicular cargo receptor and a marker of COPI coated vesicles, whereas in a control kidney and a kidney biopsy of the male with classic FD, AGAL localized to lysosomes (**Figure 4**). A similar pattern of AGAL distribution was detected in cultured skin fibroblasts (**Supplementary Figure S2**).

**Figure 4.**
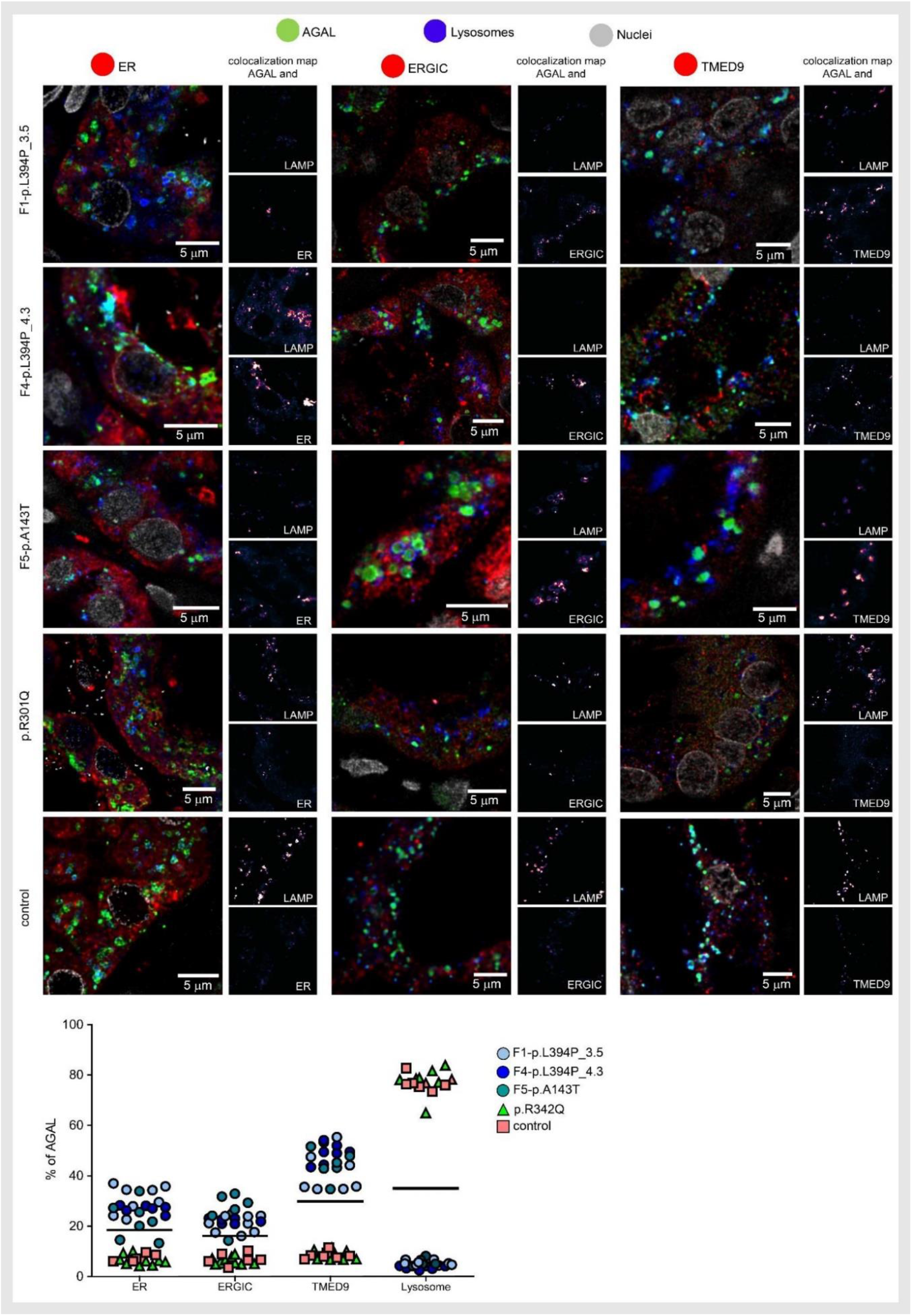
AGAL localization in kidney. To assess whether and how the identified variants affect AGAL intracellular localization in tubular epithelial cells, we detected AGAL (green) and co-localized the resulting immunofluorescent signal with markers of lysosomes (LAMP2; blue) and endoplasmic reticulum (PDI; red), endoplasmic reticulum Golgi intermediate compartment (ERGIC53; red) and the secretory cargo receptor TMED9 (red) in kidney biopsies. Nuclei are in gray. In kidney tissue from the male with the p.L394P variant (F1-394P_3.5 and F4-394P_4.3, see pedigrees), AGAL localizes mainly to ER, ERGIC and co-localizes with TMED9 with minimal staining of the lysosome. A similar pattern of AGAL distribution was found in the kidney from a female with a p.Al43T variant (F5-143T). In a kidney biopsy of a male with classic FD and p.R301Q pathogenic variant and in a control kidney, AGAL localized mostly to lysosomes with minimal staining in other compartments. The degree of AGAL colocalization with selected markers is demonstrated by the fluorescent signal overlap coefficient values that ranges from 0-1. The resulting overlap coefficient values are presented as the pseudo color whose scale is shown in the corresponding lookup tables (LUT). Subcellular distribution of AGAL in selected compartments are shown in the graph below. The values represent the percentage of the total of mean AGAL colocalization signal intensities in corresponding compartments. On average, 50 cells were analysed and 5000-15000 events were identified for each sample.

Given the clinical presentation, histopathologic, ultrastructural and fibroblast culture findings, we reasoned that misprocessing with abnormal intracellular distribution of the p.L394P AGAL might disrupt the secretory pathway, activate the UPR and cause kidney disease, similar to several other protein-misfolding disorders [21–24]. At the same time, a portion of the mutated AGAL might escape secretory pathway quality control mechanisms, transit to lysosomes, and its residual enzymatic activity might prevent intracellular accumulation of Gb3Cer and related glycosphingolipids that is characteristic in FD.

Accordingly, *in silico* analysis revealed that the L394 residue is located outside of the AGAL active site and suggested that the enzymatic activity is structurally maintained while protein misfolding is likely to occur (**Supplementary Figure S3).** Consistent with immunofluorescent studies in human kidney and cultured skin fibroblasts, the p.L394P AGAL with a C-terminal FLAG tag (p.394P AGAL-FLAG) stably expressed in HEK293 cells localized mainly to the ER and ERGIC, with preferential codistribution with the TMED9, and with minor presence in lysosomes. The wild-type AGAL with a C-terminal FLAG tag (wt AGAL-FLAG) and classic FD causing variant p.R301Q AGAL with a C-terminal FLAG tag (p.301Q AGAL-FLAG) localized mainly to lysosomes (**Figure 5**). A small proportion of all the AGAL variants localized to the Golgi apparatus (data not shown). RNA sequencing and proteomic analyses of HEK 293 cells stably expressing the p.394P AGAL-FLAG revealed activation of ER stress and UPR (**Supplementary Table S2**), with specific upregulation of the activating transcription factor 6 (ATF6) and general (Complex) branches. This response was absent in HEK 293 cells stably expressing the wt AGAL-FLAG or classic FD causing variant p.112H AGAL-FLAG. The proteins involved in the ribonuclease inositol-requiring protein-1 (IRE1) and the ER resident transmembrane protein kinase (PERK) pathways showed minimal changes (**Figure 5**).

**Figure 5.**
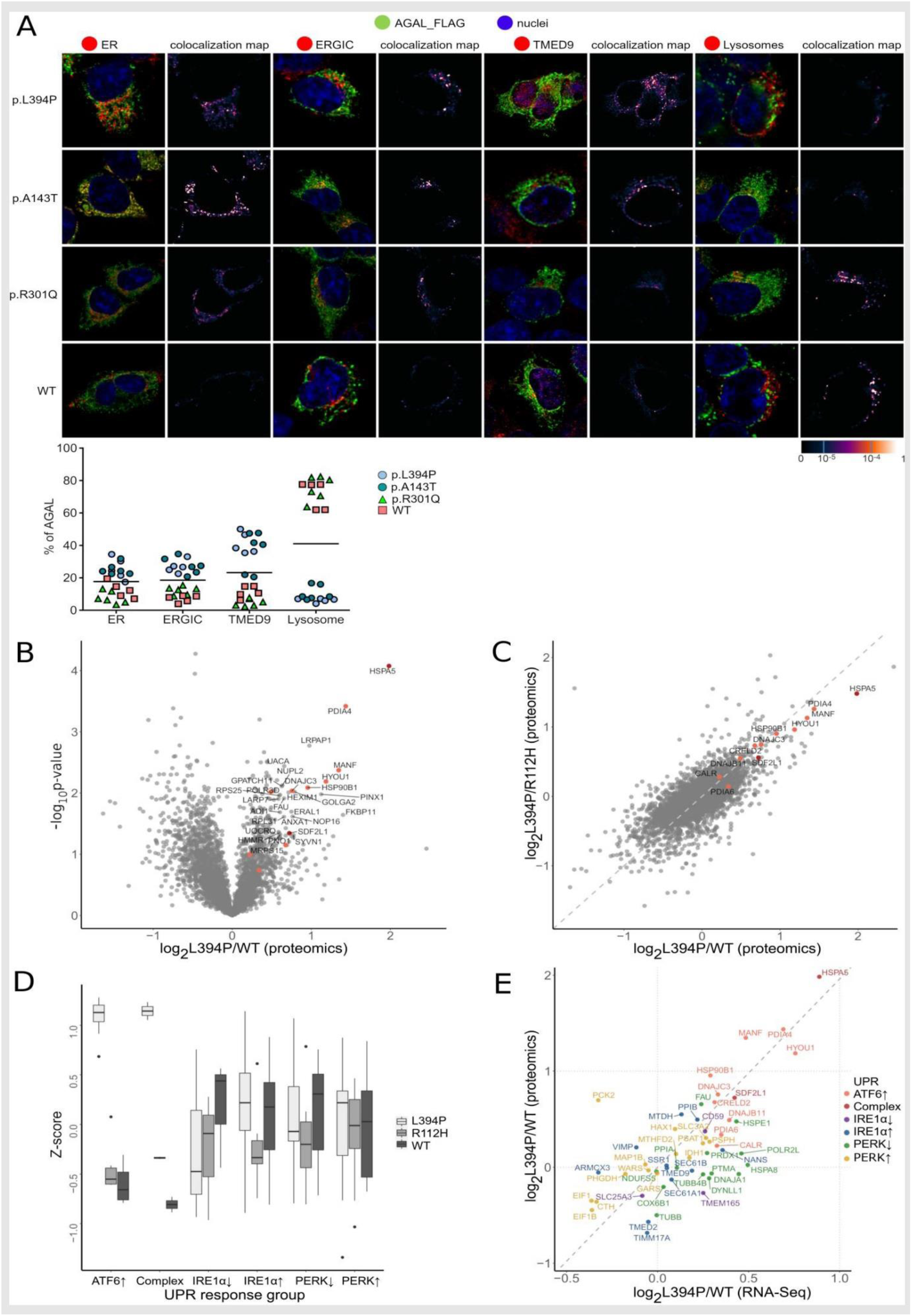
Pathogenic consequences of the p.L394P AGAL. Consistent with kidney biopsies, the corresponding AGAL-FLAG proteins stably expressed in HEK 293 also had a different intracellular distribution. We detected AGAL-FLAG (green) and co-localized the resulting immunofluorescent signal with markers of endoplasmic reticulum (PDI; red), endoplasmic reticulum Golgi intermediate compartment (ERGIC53; red), the secretory cargo receptor TMED9 (red) and lysosomes (LAMP2; red). Nuclei are stained in blue. The p.394P and p.143T AGAL-FLAG proteins localized mainly to the ER, ERGIC and co-localized with TMED9 with minimal staining iof the lysosomes. The p.301Q (classic Fabry mutation) and the wild-type AGAL-FLAG proteins localized mostly to lysosomes with minimal staining in other compartments. The degree of AGAL colocalization with selected markers is demonstrated by the fluorescent signal overlap coefficient values that ranging from 0-1. Subcellular distribution of AGAL in selected compartments are shown in the graph below. The values represent the percentage of the total mean AGAL colocalization signal intensities in corresponding compartments. On average, 50 cells were analysed and 5000-15000 events were identified for each sample. (B) Protein composition of HEK 293 cells stably expressing L394P, R112H (classic Fabry) and WT AGAL proteins was analyzed using mass spectrometry. The binary logarithm of the ratios of individual protein amounts between the cells expressing p.L439P and WT proteins (log2(L394P/WT) and the decadic logarithm of the nominal probability that the protein is differentialy expressed (-log10 (raw p-value)) is visualized using a volcano plot. Proteins on the right (positive) side of the X axis are increased in the cells expressing the p.L394P AGAL. Red dots represent proteins involved in the unfolded protein response. (C) Correlation of individual protein amount ratios in HEK 293 cells stably expressing p.L394P AGAL with p.RU2H (classic Fabry) (log2(L394P/RH2H) and WT AGAL proteins (log2(L394P/WT) shows that the presence of the p.L394P AGAL protein specifically increases the amount of protein involved in the unfolded protein response (red dots). (D) Specific activation of ATF6 and Complex branches of the unfolded protein response (UPR) in HEK 293 cells expressing the p.L394P AGAL. Z scores of normalized protein expression values were obtained from proteomic analysis of HEK 293 cells stably expressing p.L394P AGAL, p.RU2H (classic Fabry) and WT proteins. (E) Correlation of protein amounts and mRNA levels of individual UPR components in HEK 293 cells expressing the p.L394P AGAL and wild-type (WT) AGAL.

The initial step in IRE1 response, the X-Box Binding Protein 1 (XBP1) mRNA splicing, was also not activated (not shown). Consistently, expression of the ER stress and ATF6 inducible protein CRELD2 [25], was induced in kidney tubular cells of the male with the p.L394P AGAL, but not of the classic FD and healthy control (**Figure 6**).

**Figure 6.**
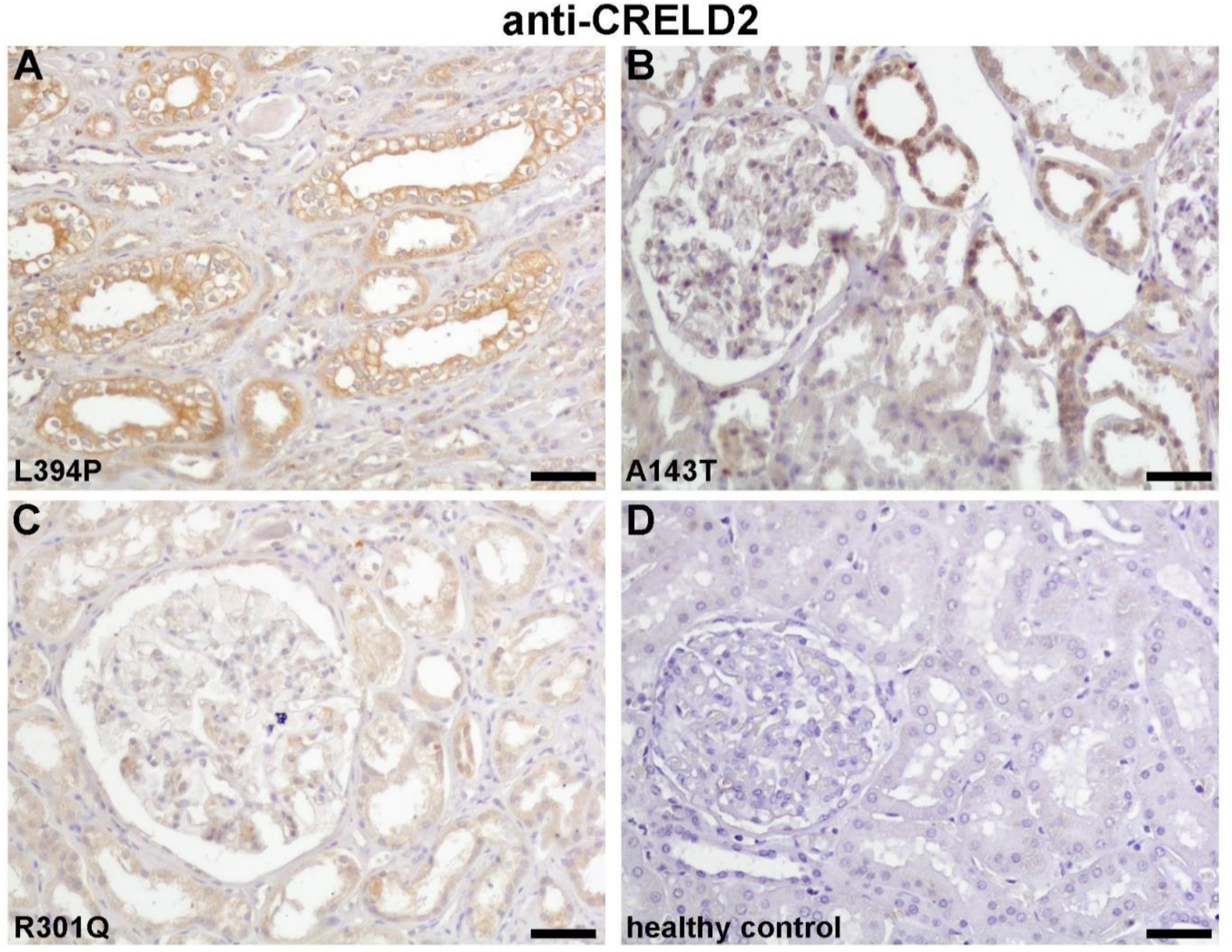
Immunohistochemical detection of the ER stress and ATF6 inducible protein CRELD2 in kidney. (A) Consistent with transcriptomic and proteomic analyses of stably transfected HEK 293 cells, the expression of CRELD2 is induced in renal tubular cells from a male with the p.L394P variant. (B) Expression of CRELD2 is induced but to a lesser extent in renal tubular cells from a female with the p.A143T variant. (C) CRELD2 expression is negligible in a male with classic FD with the p.R301Q variant. (D) CRELD2 is not expressed in control female kidney. Scale bars – 50 mm.

After identification of abnormal intracellular trafficking and UPR activation with partial slip-through of the enzymatically competent p.L394P AGAL to lysosomes, we explored similar changes in other known AGAL variants associated with FD type 2 or causing FD type 1 **(Supplementary Table S3)**. In a kidney biopsy of a female with the relatively common variant p.A143T AGAL [26], AGAL had a similarly abnormal intracellular distribution to that of patients with the p.L394P AGAL variant (**Figure 4**). Furthermore, renal tubular cells showed induction of CRELD2 expression but to a lesser extent than was detected in individuals with the p.L394P AGAL variant (**Figure 6**). Accordingly, in the HEK 293 cells stably expressing the 143T variant of AGAL with a C-terminal FLAG tag (p.143T AGAL-FLAG), AGAL localized mainly to the ERGIC with preferential co-distribution with the cargo receptor TMED9, and with a minor presence in lysosomes (**Figure 5**). Similar localization of AGAL to ERGIC and co-distribution with TMED9 that were characteristic for p.L394P and p.A143T variants, were observed also with several other nontruncating AGAL variants either of unknown significance or identified in individuals with non-classic later-onset FD, when transiently expressed in HEK 293 cells.

AGAL variants associated with classic FD always localized to lysosomes **(Supplementary Figure S4; Figure 7A**). Irrespective of their intracellular distribution and phenotypic associations, most of the tested nontruncating missense variants limited AGAL secretion. Deficit in AGAL secretion was proportional to the intensity of the ER stress and UPR activation **(Figure 7B,C**).

**Figure 7.**
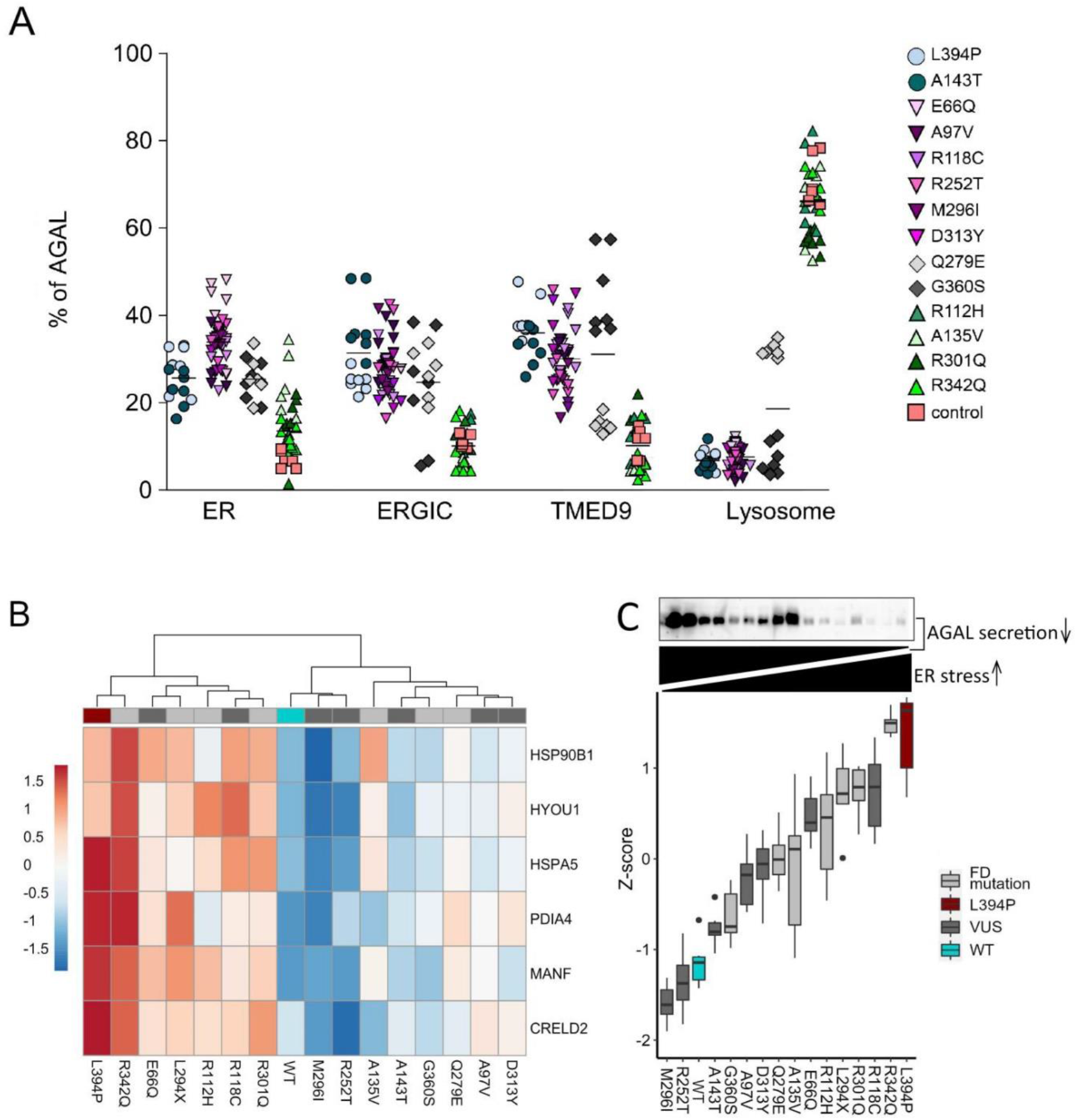
Localization of AGAL-FLAG in transiently transfected HEK 293 cells. **(A)** To assess how other nontruncating missense AGAL variants with conflicting pathogenicity affect intracellular localization, we detected corresponding AGAL-FLAG proteins transiently expressed in HEK 293 cells and co-localized the resulting immunofluorescent signal with markers of endoplasmic reticulum (PDI), endoplasmic reticulum Golgi intermediate compartment (ERGIC53), the secretory cargo receptor TMED9 and lysosome (LAMP2). Subcellular distribution of AGAL in selected compartments is shown in the graph. The values represent the percentage of the total mean AGAL colocalization signal intensities in corresponding compartments. On average, 50 cells were analysed and 5000-15000 events were identified for each sample. (B) Unsupervised hierarchical clustering of normalized expression values (Z-scores) of selected ER stress and UPR markers obtained from RT-qPCR analyses of HEK 293 cells transiently expressing individual AGAL variants. The Z-scores are presented as the pseudo-color whose scale is shown in the corresponding lookup table. The p.L394P AGAL variant is shown as a dark red; AGAL variants associated with either classic or non-classic later-onset FD are shown in light gray; AGAL variants of unknown significance (VUS) are shown in dark gray, wild type AGAL is shown in blue. (C) Intensity of the ER stress shown as Z-scores of means of normalized mRNA expression values of selected ER stress and UPR markers calculated for each of the variant. The ER stress intensity (dark) is proportional to secretory incompetence of the corresponding variant shown above by immunodetection of AGAL in cell culture media.

## Discussion

In this work we characterized the clinical, biochemical, genetic, molecular, cellular and organ pathology correlates of the p.L394P AGAL variant that was identified in six individuals with end-stage kidney disease by the Czech national screening program for FD in patients receiving renal replacement therapy. An additional twenty four carriers of this variant were identified by genetic screening of affected families. The patients presented predominantly with proteinuria, hypertension, chronic kidney disease, and heart disease. The median age of ESRD was older than that of classic FD. Characteristic findings such as angiokeratomas and neuropathy were less commonly seen. Interestingly, type 2 diabetes mellitus was detected in 40% of patients diagnosed with the p.L394P AGAL variant vs. 0% of family members who did not have this variant. Diabetes mellitus has not been previously reported with FD. This finding, only observed in the current study, could be coincidental, but it would be worthwhile to check for the prevalence of diabetes in other families with classic and non-classic FD. Women were more likely to be affected than in classic FD but had milder phenotype with later age at onset than males. Affected males presented with ~15% residual AGAL activity in plasma and leukocytes. Female carriers had normal AGAL activities. In cultured male skin fibroblasts, the residual activity of p.L394P AGAL resulted in decreased Gb3Cer degradation that was intermediate between control and classic FD fibroblasts, likely explaining normal plasma lyso-Gb3Cer levels in affected carriers. Accordingly, characteristic lysosomal storage was absent in all kidney cell types [27]. Biopsy specimen, however, showed mild interstitial fibrosis and podocytopathy (effacement of podocyte pedicels, and non-membrane bound lipid droplets in proximal tubular epithelium and interstitial cells). Interestingly, pedicel effacement was very similar to Gb3Cer- independent podocyte changes reported in the zebrafish AGAL deficient model [13] thus providing further evidence for lysosomal storage independent mechanism of FD pathogenesis.

The pathogenesis and clinical manifestations of classic FD have been primarily attributed to the absence or profound enzymatic deficiency of AGAL, accumulation of Gb3Cer and consequent disarrangement(s) of intracellular pathways linked to lysosomal (dys)function(s) [12]. Individuals with classic FD commonly carry truncating *GLA* variants resulting in loss of AGAL and/or its enzymatic activity. However, approximately 30% of reported pathogenic *GLA* variants are missense variants [9]. Many of these variants result in non-classic FD, with lesser organ involvement, later age of onset, and >5% of residual enzyme activity [5]. Moreover, many *GLA* variants of conflicting pathogenicity or unknown significance have been identified within screening programs for FD in newborns and high-risk populations [6]. In many individuals affected with these variants and presenting with atypical clinical features of FD, other factors may be contributing to disease pathogenesis besides the AGAL enzymatic deficiency and lysosomal storage.

For many missense AGAL variants associated with the classic FD, enzyme deficiency results from misfolding and endoplasmic reticulum associated protein degradation (ERAD) [28]. In amenable variants, pharmacological chaperones allow correct folding, stabilization and trafficking of mutated AGAL proteins through the secretory pathway to the lysosomes to catabolize accumulated substrates [4, 29, 30]. Instead of being degraded via ERAD or transported to the lysosome as in classic FD, most of the p.L394P AGAL localizes to ERGIC and TMED9 positive cargo receptor-containing vesicles, where the protein folding quality control system is present. The interference of misprocessed AGAL with the proper function of the ER, ERGIC, and TMED9 specifically, leads to the ER stress and activation of UPR, presumably triggering effects with pathogenic consequences on secretory active and/or post mitotic cell types. Similar dearrangements have been identified as primary in various monogenic glomerular and tubular diseases [24] endothelial dysfunction [31], cardiometabolic diseases [32], type 2 diabetes [33] and neurodegeneration [34].

Stimulated by the finding of potential ERGIC/TMED9 involvement and UPR activation in the kidney biopsies of the p.L394P AGAL carriers, we investigated several other clinically described *GLA* variants. First, we studied a female from our cohort with the p.A143T AGAL, the most common variant in the US, and the genetic cause of the initial case of FD reported by William Anderson [35]. She had normal AGAL activity, plasma lyso-Gb3Cer and urinary Gb3Cer levels and presented with late onset chronic kidney disease. Her kidney biopsy showed mild interstitial fibrosis and podocytopathy with absence of lysosomal storage. Intracellular AGAL distribution and signs of UPR activation were very similar to the p.L394P AGAL carriers. The p.A143T variant, which we found to be associated with ERGIC deposition, has been considered nonpathogenic in several case series studies [6, 26]. However, other studies consider the p.A143T variant to be a cause of non-classic FD with incomplete age-related and sex-related penetrance and predominantly cardiac manifestations [36].

We then studied intracellular distribution of several other missense AGAL variants in either stably or transiently transfected HEK 293 cells. Pathogenic variants associated with classic and non-classic FD (p.R112H, p.A135V, p.L294X, p.R301Q and p.R342Q) localized to lysosomes and demonstrated characteristic biochemical abnormalities and histopathologic findings in kidney biopsies. Representative missense AGAL variants associated either with non-classic later-onset FD (p.A97V, p.Q279E and p.M296I) or variants of unknown significance (p.L394P, p.E66Q, p.R118C, p.A143T, p.R252T and p.D313Y) localized to the ER and ERGIC.

Most of the tested missense variants limited AGAL secretion. The intracellular retention of AGAL always activated ER stress and UPR in affected tissues or transfected cells. The intensity of ER stress and UPR varied and was proportional to the secretory deficit of the corresponding AGAL variant. There were no differences in ER stress and UPR activation between missense AGAL variants retained along the secretory pathway and associated with non-classic FD (like the p.L394P) and AGAL variants localized to the lysosome and causing the classic FD (like the p.R342Q). Thus AGALopathy seems to be an important component in the pathogenesis of FD, independent of the enzymatic deficiency and lysosomal storage.

As with many genetic diseases, genotype-phenotype correlations are difficult and carrying a specific variant does not necessarily mean that disease will develop. It was impossible to correlate our genetic and molecular findings with clinical findings in the literature because of the small number of individuals reported with each mutation and the non-systematic reporting of clinical features.

We believe that the underlying pathogenesis of FD symptoms in carriers of missense AGAL variants is related to a combination of decreased enzymatic activity, altered lipid composition of membranes due to limited Gb3Cer processing, intracellular AGAL retention, chronic ER stress and UPR activation. The extent of each factor may be determined by the type of the variant and host factors, with each factor contributing more or less to the clinical findings (**Figure 8**).

**Figure 8.**
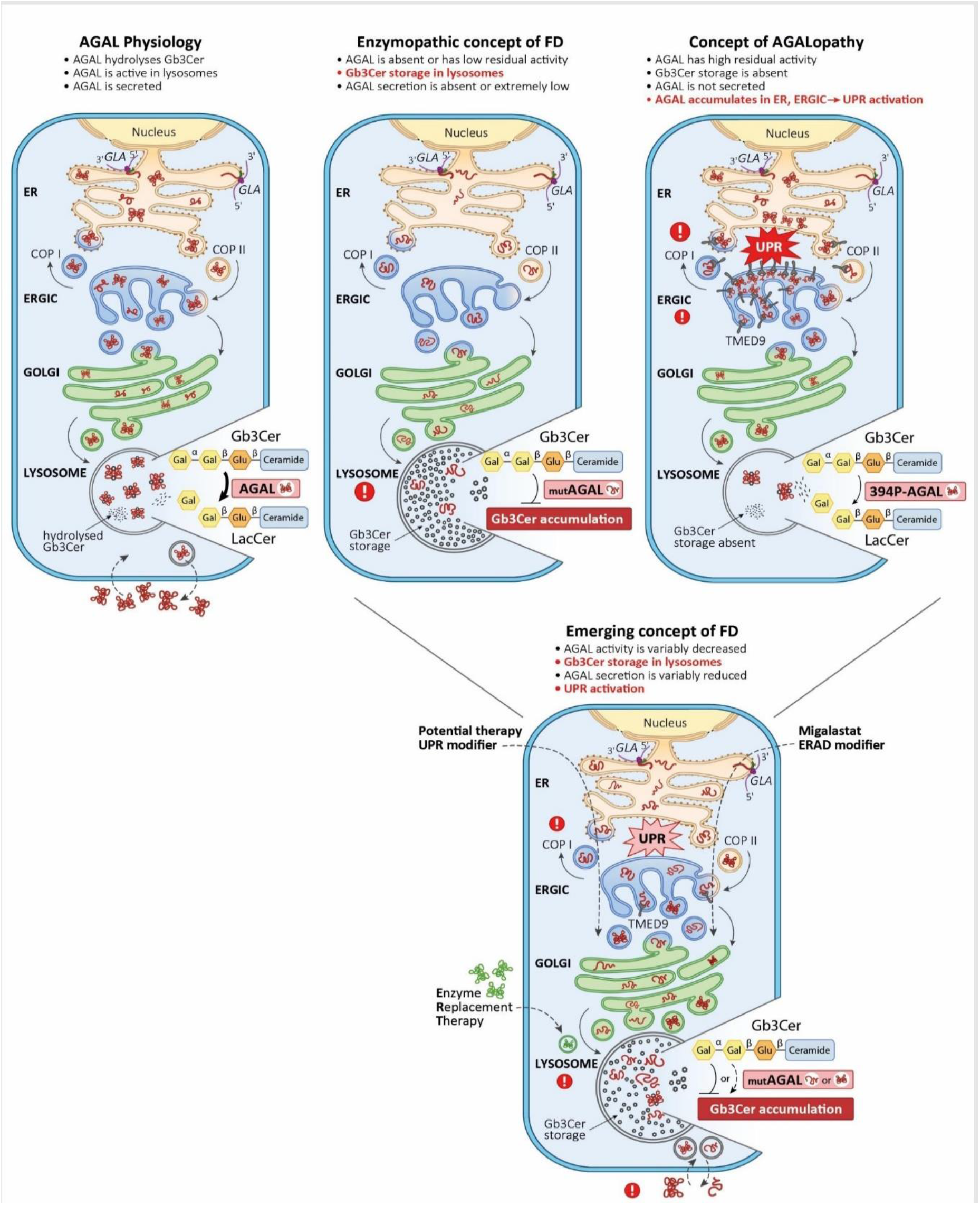
ER stress and unfolded protein response in Fabry disease. AGAL is **a** homodimeric glycoprotein that hydrolyzes the terminal alpha-galactosyl (α-Gal) moieties from glycolipids and glycoproteins in lysosomes. During biosynthesis *GLA* mRNA is co-translationally translocated into the endoplasmic reticulum (ER).This process is initiated in the cytoplasm and mediated by the signal peptide (green). In the ER, nascent AGAL undergoes a series of post- translational modifications, including removal of the signal peptide, N-glycosylation, and chaperone-mediated folding. Properly folded AGAL exits the ER in the coat protein complex II (COPII)-coated vesicles and enters the endoplasmic reticulum Golgi intermediate compartment (ERGIC), where concentration, folding, quality control and sorting of newly synthesized proteins occurs. Unmaturated proteins are recycled back to the ER via COPI-coated vesicles. Mature AGAL traverses through Golgi to lysosomes, where it is activated and hydrolyses the terminal alpha-galactosyl moietiesfrom glycolipids and glycoproteins. A proportion of AGAL is secreted and re-endocytosed. The “classic” enzymopathic concept of Fabry disease (middle panel) claims that *GLA* variants leading to the loss of AGAL protein or of its enzymatic activity lead to deposition of globotriaosylceramide (and other glycoshingolipids) in lysosomes. Clinical manifestations are believed to develop as a result of the storage-induced dysfunction of lysomal and lysosomal-associated cellular compartments and signaling pathways. AGAL secretion is virtually absent. AGALopathic concept is evidenced by the pathogenic impacts of the p.L394P AGAL variant. Misfolding and abnormal intracellular accumulation of this missense AGAL variant in ER and ERGIC disrupt the early secretory pathway and activate the unfolded protein response (UPR). At the same time, a portion of the mutated but enzymatically competent AGAL escapes secretory pathway quality control mechanisms and transits to lysosomes where it prevents accumulation of globotriaosylceramide (Gb3Cer) and related glycosphingolipids. L394P AGAL is not secreted. Similar effects (accumulation in ER and ERGIC, UPR activation) can be variably induced also by other missense *GLA* mutations. Secretion of such AGAL variants is decreased proportionally to their defective intracellular trafficking. We believe that the enzymopathic and AGALopathic components should be considered as key contributors to the pathogenesis of Fabry disease. We further propose that both these components contribute to overall pathogenic impacts of individual AGAL variants. Some variants have more enzymopathic and some more AGALopathic nature. Unlike p.L394P AGAL, which is predominantly AGALopathic, most of the *GLA* missense mutations trigger mixed (enzymopathic and AGALopathic) downstream effects. Whereas FD patients with predominantly enzymopathic variants are likely to respond well to ERT or chaperone therapy targeting ERAD, individuals with preferentially AGALopathic variants may be adversely affected by defective trafficking of the misfolded AGAL and augmented consequences of intracellular stress and UPR. For these variants, compounds modulating UPR may provide new therapeutic opportunities.

In this investigation, we also found that females with the p.L394P AGAL variant were more likely to be affected than in classic FD but had milder phenotype with later age at onset than males. A similar clinical course has also been seen in other females with non-classic FD, often in families with missense variants [37]. This finding is consistent with the concept of mosaic AGALopathy of the p.L394P AGAL and similar missense AGAL variants rather than loss of enzymatic function of one allele. Moreover, AGALopathy may explain why females affected by FD become symptomatic in the setting of normal or mildly decreased AGAL activity and why the wild-type AGAL secreted from cells expressing the wild type *GLA* allele does not correct ER stress and UPR within cells expressing the mutated one.

The mechanism of AGALopathy being a lysosomal storage independent etiologic factor in FD could also explain why some patients with missense variants treated with recombinant AGAL do not respond to enzyme replacement therapy [3]. AGALopathy may also explain, why some patients treated with Migalastat experienced accelerated loss of kidney function [38]. This intervention may adversely enhance transport of the misfolded AGAL from ER to the ERGIC, where it will further increase negative consequences of intracellular stress and UPR. Moreover, the pathogenic concept of AGALopathy resulting from protein misprocessing in the secretory pathway may also apply to phenotypic variabiality in other lysosomal storage diseases with high residual enzymatic activity(ies) [39].

In conclusion, our results suggest that aberrant intracellular trafficking of some missense AGAL and pathogenic interaction within the ERGIC/TMED9 leading to maladaptive ER stress response may explain some of the varied pathogenic and clinical findings in FD. Recent research identifying compounds modulating UPR [40], and facilitating removal of unfolded proteins from TMED9 enriched transport vesicles into lysosomes [22], provide new opportunities to probe the pathologic and potentially therapeutic implications in FD.

## Methods

This investigation was approved by the Ethics Committee of the First Faculty of Medicine of the Charles University, Prague, Czech Republic, and all participants provided informed consent.

### Patients

The probands were identified through a nationwide Fabry disease screening program among patients receiving maintenance dialysis therapy in the Czech Republic. This included all dialysis units (N=112) operating in the Czech Republic in the screening period between September 1, 2016 to May 30, 2018, and thus finally sampling and testing altogether 6352 patients (i.e. 93.9% of prevalent patients registered as chronic dialysis patients in 31.12.2017). After obtaining informed consent, peripheral blood was obtained for measurements of AGAL enzymatic activity, globotriaosylsphingosine (lyso- Gb3Cer) concentration and *GLA* sequencing at CentoCard, CentoGene AG.

### Biochemical analyses

AGAL activity in plasma and leukocytes was measured by a fluorometric method using 4- methylumbelliferyl-α-D-galactopyranoside as the substrate in the presence of N-acetylgalactosamine as an inhibitor of α-D-galactosidase B [41]. Serum creatinine and other biochemical parameters were measured using routine clinical methods.

### LC-MS analysis of lyso-Gb3Cer, Gb3Cer and met-Gb3Cer

Plasma lyso-Gb3Cer was extracted using the modified method of Aerts et al [42]; urinary Gb3Cer and met-Gb3Cer were extracted using the modified method of Abaoui et al [43]. The samples were analyzed on AB/MDS SCIEX API4000 triple quadrupole tandem mass spectrometer with electrospray ionization coupled to the Agilent 1290 Infinity UPLC system. The mass spectrometry was performed in positive ion mode and SRM using [M+H^+^]^+^ precursor ions later fragmented via CID using nitrogen as collision gas. The product ions with m/z of 282 (lyso-Gb3Cer), 289 (D7-lyso-Gb3Cer IST) and corresponding ceramide for Gb3Cer and met-Gb3Cer were used for quantitative analysis. The quantity was calculated using external calibrator consisting of 25ng of D7-lyso-Gb3Cer or (D3-C18:0, d18:1) Gb3Cer combined with 50ng of lyso-Gb3Cer or Gb3Cer according to quantified lipid as previously used [44].

### Degradation of globotriaosylceramide in cultured skin fibroblasts

Skin fibroblasts were loaded with mass labeled C23:0,d18:1 globotriaosylceramide (C23Gb3Cer). Conduritol B epoxide (Calbiochem-Novabiochem GmbH, Germany), a covalent inhibitor of GlcCer-β- Glucosidase, was added to block the downstream metabolic conversion of glucosylceramide to ceramide. After 4 days, the medium was removed, cells were harvested and sphingolipids were extracted from the cell homogenate and quantified using the internal standard C17:0 globotriaosylceramide and the flow injection analysis electrospray ionization tandem mass spectrometry (FIA-ESI-MS/MS) on the AB/MDS SCIEX API4000 triple quadrupole tandem mass spectrometer coupled to UPLC Agilent 1290 Infinity as previously described [44, 45].

### Genetic evaluation

Genomic DNA of all available individuals was extracted from whole blood in a standard manner. The *GLA* variant (chrX:100652906 A>G (hg19; c. 1181T>C) was determined using Sanger sequencing. Whole genome sequencing, data analysis, and identity-by-descent mapping was performed as described [46].

### Histopathological, immunohistochemical and ultrastructural electron microscopic analyses of kidney biopsies

Paraformaldehyde-fixed kidney biopsies of family members **F1_L394P_V.5** and **F4_L394P_IV.3** were investigated. AGAL was detected with rabbit monoclonal anti-galactosidase alpha antibody (Abcam, ab168341, Cambridge, UK); CRELD2 was detected with mouse polyclonal anti-CRELD2 antibody (Sigma-Aldrich, SAB14001820, Prague, Czech Republic); lysosomes were detected with rabbit polyclonal anti-human LAMP-1 antibody (kindely provided by dr. Carlsson, University of Umea, Sweden). Detection of bound primary antibody was achieved using Dako EnVision + TM Peroxidase Rabbit Kit (DAKO, Glostrup, Denmark) or Dako EnVision + TM Peroxidase Mouse Kit (DAKO, Glostrup, Denmark) with 3,3’-diaminobenzidine as substrate. Ultrastructural electron microscopic studies were performed as reported earlier [47].

### Intracellular localization of AGAL in kidney biopsies

For intracellular localization of AGAL in kidney, we stained always in parallel AGAL and LAMP-2, a marker of lysosomes, together with markers of the ER, ERGIC, Golgi and COPI coated vesicles. AGAL was detected with the rabbit monoclonal anti-galactosidase alpha antibody (Abcam, ab168341, Cambridge, UK); lysosomes were detected with goat polyclonal anti-human LAMP-2/CD107b antibody (R and D systems, AF6228, Minneapolis, USA); ER was detected with mouse anti-PDI (ADI-SPA-891, Enzo, Farmingdale, NY, USA); ERGIC was detected with mouse monoclonal anti-LMAN1 antibody (clone OTI1C5, Invitrogen, Thermo Fisher Scientific, Waltham, MA, USA); Golgi apparatus was detected with mouse anti-Golgi 58K protein /formiminotransferase cyclodeaminase (FTCD) antibody (G2404, SIGMA-Aldrich, Prague, Czech Republic). The following secondary antibodies were used for detection: donkey anti-rabbit IgG Alexa Fluor 555, donkey anti-goat IgG Alexa Fluor 488 or donkey anti-mouse IgG Alexa Fluor 647 secondary antibodies (Thermo Fischer Scientific). Colocalization of AGAL, lysosome and COPI coated vesicles was achieved consequently, first with AGAL and lysosome detected as described above and then with subsequent detection of TMED9 with rabbit polyclonal anti-TMED9 antibody (21620-1-AP, Proteintech, Manchester, UK) directly labeled with Daylight 488 Fast Conjugation Kit (Abcam, Cambridge, UK).

### Intracellular localization of AGAL in primary skin fibroblasts

For intracellular localization of AGAL in primary skin fibroblasts, we stained AGAL with individual markers of lysosomes, ER, ERGIC and COPI coated vesicles. AGAL, ER and ERGIC were detected as described above; lysosomes were detected with mouse monoclonal anti-LAMP2 antibody (Ab25631, Abcam); Detection of bound primary antibodies were achieved with donkey anti-mouse IgG Alexa Fluor 488 and donkey anti-rabbit IgG Alexa Fluor 555 secondary antibodies (Thermo Fischer Scientific). Colocalization of AGAL and COPI coated vesicles was achieved as described above.

### Intracellular localization of AGAL in HEK 293 cells stably or transiently transfected with *GLA_FLAG cDNA*

Wild-type *GLA* cDNA (including 5’-UTR) was reverse transcribed from human leukocyte total RNA, polymerase chain reaction (PCR) amplified, cloned into pCR- XL-TOPO vector and introduced into the One Shot^®^ TOP10 Chemically Competent *E. coli* (Invitrogen, Paisley, UK). A single 3-terminal F*LAG*-tag sequence was appended to the originally synthesized wild type *GLA* cDNA using PCR with specific oligonucleotide primers containing the BamHI and Xho I restriction sites for subsequent subcloning of the *GLA_FLAG* construct to pCR3.1 vector (Invitrogen, Paisley, UK). The resulting product *GLA_FLAG/pCR3.1* was introduced into the Escherichia coli TOP 10’F strain and clones with the correct *GLA* sequence were selected by Sanger sequencing. Mutant constructs (mut_*GLA_FLAG*/pCR3.1) were prepared by site-directed mutagenesis (Site-Directed Mutagenesis Kits, QuickChange II, Agilent) and selected by sequencing.

For stable AGAL-FLAG expression, HEK 293 cells were transfected with 2.5 μg of plasmid DNA using Lipofectamine 3000^TM^ (Invitrogen, Paisley, UK). Three days post transfection, cells were trypsinised, diluted and cultured in selective medium containing 0.5 mg/ml G418 (Invitrogen-Gibco, Paisley, UK). For each selected clone, the presence and correct sequence of *GLA_FLAG* was confirmed by Sanger sequencing; amount of *GLA*_*FLAG* transcript was determined by real-time quantitative PCR, and expression levels of AGAL-FLAG tagged proteins were assessed by Western blot with mouse monoclonal anti-FLAG antibody (F1804, SIGMA-Aldrich, Prague, Czech Republic). Clones demonstrating similar *GLA_FLAG* transcript and AGAL-FLAG protein amounts were selected for further analyses.

For transient AGAL-FLAG expression, transfections were carried out using Lipofectamine 3000^TM^ (Invitrogen, Paisley, UK) with either 0.5 μg or 2.5μg DNA for 1.5×10^5^ or 8×10^5^ cells, respectively. AGAL-FLAG was detected 24 hours post transfection by Western blot analysis with mouse monoclonal anti-FLAG antibody (F1804, SIGMA-Aldrich, Prague, Czech Republic). Endogenously expressed AGAL and tubulin were detected with rabbit monoclonal anti-galactosidase alpha antibody (used for staining of AGAL in kidney) and mouse monoclonal anti-acetylated tubulin antibody (T7451, SIGMA-Aldrich, Prague, Czech Republic), respectively.

For intracellular localization of AGAL-FLAG in stably or transiently transfected HEK 293 cells we stained AGAL-FLAG with individual markers of lysosomes, ER, ERGIC, Golgi and COPI coated vesicles. The AGAL-FLAG was detected with mouse monoclonal Anti-FLAG M2 antibody (F1804, Sigma Aldrich); lysosomes were detected with rabbit polyclonal anti-LAMP2 (custom prepared); the ER was detected with rabbit polyclonal anti-SEC61A1 antibody (ab183046, Abcam); the ERGIC was detected with rabbit polyclonal anti-LMAN1 (ProteinTech); the Golgi apparatus was detected with rabbit monoclonal anti-GM130 (C-terminal, G7295, SIGMA-Aldrich, Prague, Czech Republic). Detection of bound primary antibodies was achieved using donkey anti-mouse IgG Alexa Fluor 488 donkey anti-rabbit IgG Alexa Fluor 555 secondary antibodies (Thermo Fischer Scientific). Colocalization of AGAL and COPI coated vesicles was achieved as described above.

### Confocal microscopy, fluorescence image acquisition and analysis

Slides were mounted in ProLong gold antifade mountant with DAPI (Thermo Fischer Scientific) and analyzed by confocal microscopy. XYZ images were sampled according to Nyquist criterion using a Leica SP8X laser scanning confocal microscope, HC PL Apo objective (63X, N.A.1.40), 405 nm diode/50mW DMOD Flexibl, with 488, 555 and 647 laser lines in 470nm-670nm 80MHz pulse continuum WLL2. Images were restored using a classic maximum likelihood restoration algorithm in the Huygens Professional Software (SVI, Hilversum, The Netherlands). The colocalization maps employing single pixel overlap coefficient values ranging from 0-1, were created in the Huygens Professional Software. The resulting overlap coefficient values are presented as the pseudo color, the scale of which is shown in the corresponding lookup tables (LUT).

Relative subcellular distribution of AGAL was calculated from the mean of AGAL signal intensities that were retrieved from pixels with positive colocalization signals by the Analyze Particles tool in ImageJ Software. On average for each sample, the signals were obtained from 50 cells and 5000-15000 events were identified.

### AGAL secretion

Secreted AGAL was detected in media of transiently transfected HEK 293 cells (see above). Media were processed as described before [23], proteins were transferred Immobilon-E PVDF membrane (Merck Millipore, Tullagreen, Ireland) and AGAL was detected by monoclonal anti-Galactosidase alpha antibody (# ab168341, Abcam, Cambridge, UK) and by HRP-conjugated goat anti-rabbit IgG (H+L) secondary antibody (# 31460, Thermo Fisher Scientific, Prague, Czech Republic)

### RNA sequencing of HEK 293 cells stably expressing AGAL-FLAG

Total RNA was isolated from stably transfected cells using the RNA Minikit (Qiagen). RNA concentrations were determined spectrophotometrically at A260 nm by NanoDrop (NanoDrop Technologies, Wilmington, DE), and RNA quality was verified using an Agilent 2100 bioanalyser - RNA Lab-On-a-Chip (Agilent Technologies, Santa Clara, USA). Stranded mRNA-Seq libraries were prepared with KAPA mRNA HyperPrep Kit for Illumina^®^ Platforms (Roche). Paired-ends reads of 2×100 base pairs were sequenced on Illumina NoveSeq6000 at the National Center for Medical Genomics in Prague according to manufacturer protocol. The resulting FASTQ files were subjected to QC control and trimmed using Atropos (version1.128) [48]. Gene-level abundances were estimated using Salmon (version 1.3) [49] with the Ensembl gene definition version75. Normalization and differential expression analyses were performed within DESeq2 R package [50].

### Proteomic analysis of HEK 293 cells stably expressing AGAL-FLAG

Cell pellet homogenates were trypsin digested, and the resulting peptides were analyzed with nano-scale liquid chromatographic tandem mass spectrometry (nLC-MS/MS) using the Thermo Orbitrap Fusion (Q-OT- qIT, Thermo Scientific). All data were analyzed and quantified with the MaxQuant software [51] (version 1.6.3.4). Quantifications were performed with the label-free algorithm in MaxQuant [52]. Data analysis was performed using Perseus 1.6.1.3 software [53]. Pathway analysis of the proteomic and RNASeq data was performed with GSEA [54] (version 4.1.0). Differentially expressed proteins were categorized on two different levels, canonical pathways (CP) and gene ontology (GO) terms (C2 and C5 gene set collections in MSigDB (version 7.2). The terms with FDR q-value <0.05 were visualized using Enrichment map version 3.3.3 plugin in Cytoscape [55] version 3.8.2. UPR branch activation analysis is presented as boxplots. Each box of the boxplot consists of proteins comprising an individual UPR branch as defined [56]. The expression of each protein in the group is plotted as mean of the scaled expression profiles obtained from three replicates as described [22].

### qRT-PCR analysis of UPR markers in HEK 293 cells transiently transfected with *GLA_FLAG cDNA*

RNA was prepared from transiently transfected HEK 293 cells using RNeasy Mini Kit (Qiagen) or TRIzol reagent (Invitrogen) according to the manufacturer’s protocol. Samples of 15 ng total RNA were reverse transcribed and amplified using SOLIScript 1-step Multiplex Probe Kit (Solis Biodyne) according to the manufacturer’s instructions. The PCR master mix contained 300 nM of each primer and TaqMan probe (Generi Biotech) in a total reaction volume of 20 μl. The qPCR reactions were run on a CFX96 instrument (Bio-Rad). Relative expression levels were calculated by 2^-ΔΔCt^ method using *PUM1, CASC3, UBC* and *TBP* genes as a reference. Primer and probe sequences used are listed in **Supplementary Table S3**.

Detailed protocols for all methods are provided in the **Supplementary Appendix**.

## Acknowledgements

We thank dr. Pešková (České Budějovice), dr. Ryba (Liberec), dr. Němcová (Jihlava), dr. Satu Pešičková, dr. Švára, dr. Ryšavá, dr. Kodet, dr. Lacina, dr. Dubská, s.n. Křečková (Prague) for their help in collecting clinical data and biologic specimens.

## Author Contributions

M. Živná and S.Kmoch conceived the concept and design of the study. M. Živná, V. Barešová, D. Mušálková, G. Dostálová, K. Hodaňová, L. Kuchař, P. Vyleťal, H. Hůlková, H. Hartmannová, V. Stránecký, A. Hnízda, M. Radina, P. Reková, L. Roblová, E. Honsová, I. Rychlík, A. Linhart and J. Sikora were responsible for acquisition and analysis of data. B. Asfaw, H. Poupětová, H. Vlášková, T. Kmochová, M. Votruba, J. Sovová, H. Trešlová, L. Stolnaja, and L. Steiner-Mrázová provided technical support. M. Živná,V. Barešová, D. Mušálková, L. Kuchař, P. Vyleťal, H. Hůlková, A. Linhart, AJ. Bleyer, J. Sikora and S. Kmoch were responsible for data interpretation. M. Živná, A. Linhart, AJ. Bleyer, J. Sikora and S. Kmoch drafted the manuscript. All authors contributed to revision of the final version of the manuscript and approved the final submitted version.

## Data Sharing Statement

The mass spectrometry proteomics data have been deposited to the ProteomeXchange Consortium via the PRIDE [57] partner repository with the dataset identifier PXD033936 and 10.6019/PXD033936.

