## Supplemental material for "AGAL misprocessing-induced ER stress and the unfolded protein response: lysosomal storage-independent mechanism of Fabry disease pathogenesis?"

This appendix has been provided by the authors to give readers additional information about their work.

**Table of contents**

**Supplementary Figure S1. (p. 2)**

Anti-LAMP1 immunohistochemistry in kidney biopsies

**Supplementary Figure S2. (p. 3)**

AGAL localization in skin fibroblasts

**Supplementary Figure S3. (p. 4)**

Structural topology and impact of the mutations

**Supplementary Figure S4. (p. 5)**

Localization of AGAL transiently expressed in HEK293 cells

**Supplementary methods and references (p. 6-14)**

**Supplementary Table S1** (*attached as a* separate document)

Clinical presentation and biochemical characteristics of individuals with p. L394P mutation

**Supplementary Table S2** (*attached as* *a* separate document)

Functional annotation of proteomic data

**Supplementary Table S3 (p. 15-18)**

The list of variants and references

**Supplementary Figure S1 Anti-LAMP1 immunohistochemistry in kidney biopsies.**

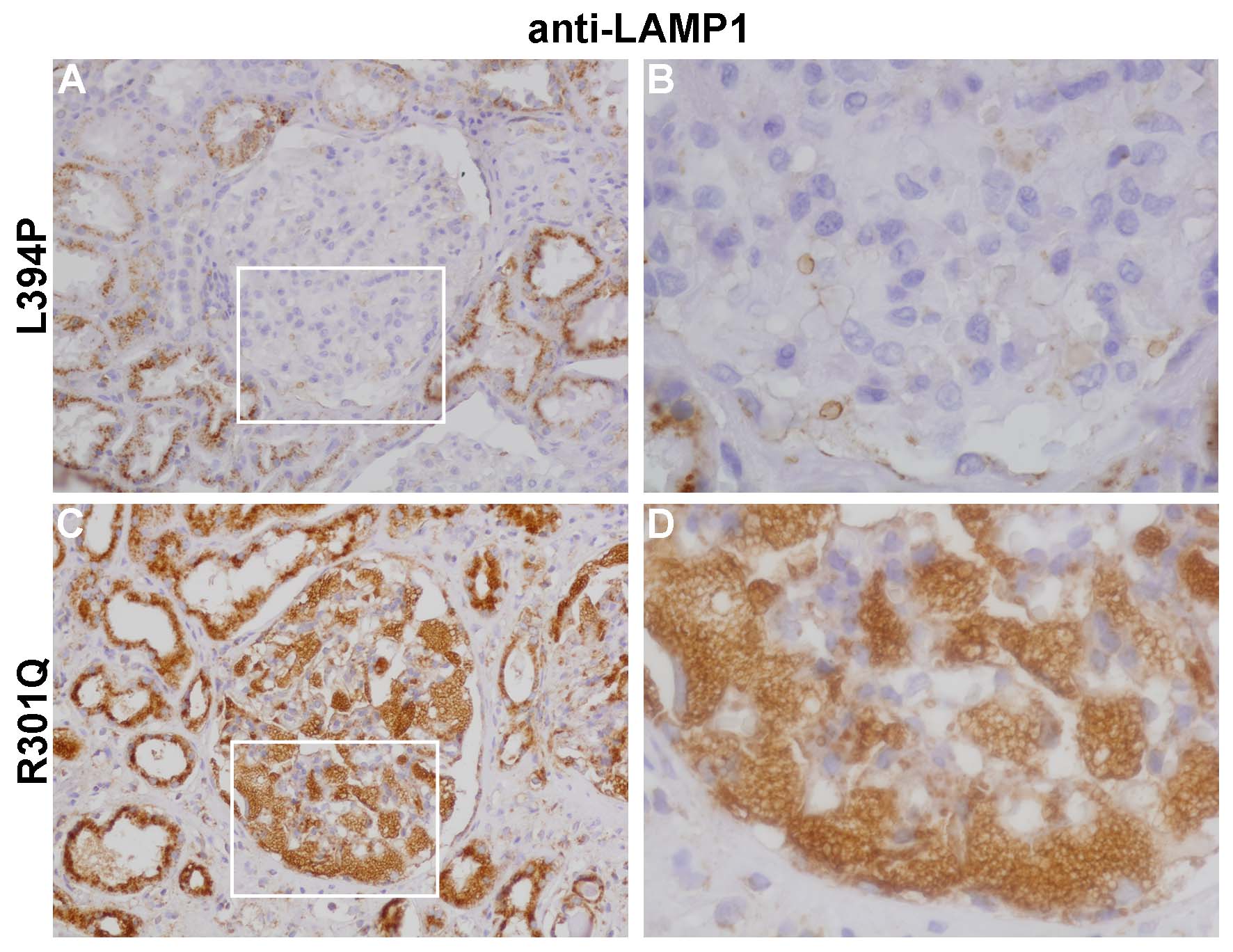

(A, B) Lysosomal storage pathology is absent in podocytes of patients with the p.L394P AGAL variant (C, D) Lysosomal storage in podocytes of a patient carrying the p.R301Q AGAL mutation. White rectangles in A and C correspond to areas shown in B and D, respectively. Scale bars = 50 μm

**Supplementary Figure S2** **AGAL localization in skin fibroblasts**

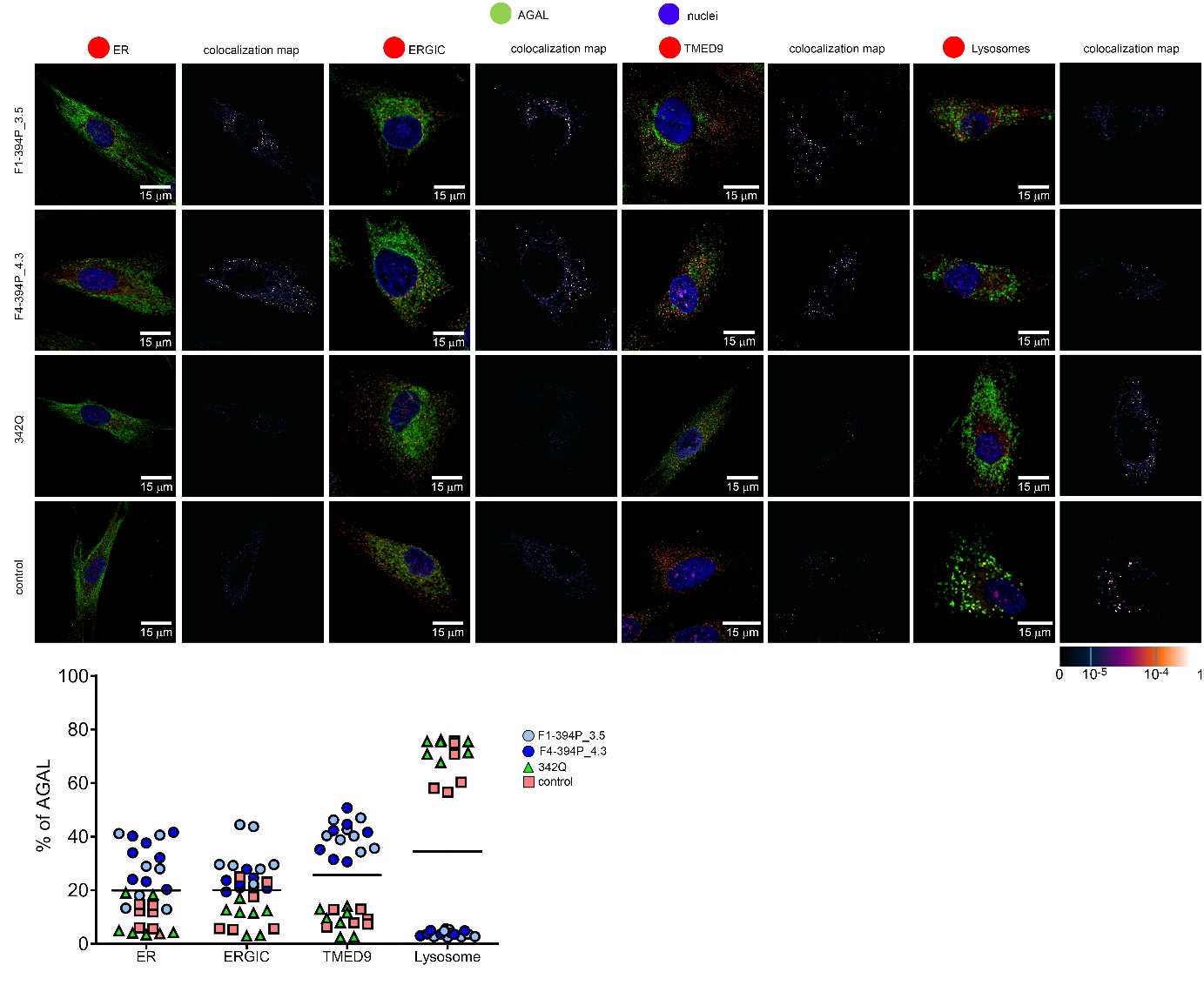

To assess whether and how identified AGAL mutations affect intracellular localization, we detected AGAL (green) and co-localized the resulting immunofluorescent signal with markers for lysosomes (LAMP2; blue) and endoplasmic reticulum (PDI; red), endoplasmic reticulum Golgi intermediate compartment (ERGIC53; red) and the secretory cargo receptor TMED9 (red) in cultured skin fibroblasts. Nuclei are in blue. In affected skin fibroblasts from individuals with the p. L394P mutation (F1-394P_3.5 and F4-394P_4.3, see pedigrees), AGAL localizes mainly to ER, ERGIC and co-localizes with TMED9 with minimal staining of the lysosome. In skin fibroblasts from a classic FD patient with the p.342Q mutation and from an unaffected control, AGAL localized mostly to lysosomes with minimal staining in other compartments. The degree of AGAL colocalization with selected markers is demonstrated by the fluorescent signal overlap coefficient values that ranges from 0-1. The resulting overlap coefficient values are presented as the pseudo color whose scale is shown in the corresponding lookup tables (LUT). Subcellular distribution of AGAL in selected compartments are shown in the graph below. The values represent the percentage of the total of mean AGAL colocalization signal intensities in corresponding compartments. On average, 50 cells were analyzed and 5000-15000 events were identified for each sample.

**Supplementary Figure S3 Structural topology and impact of the mutations in alpha-galactosidase.**

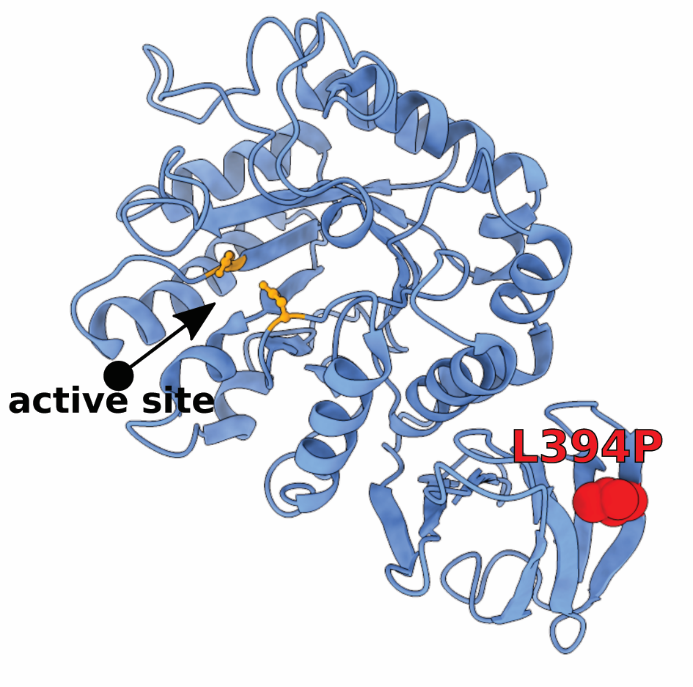

The crystal structure of AGAL (PDB ID 1R47) was used for modelling in UCSF ChimeraX. Only one subunit in homodimeric protein is shown for clarity. The active site residues are highlighted as orange sticks. Position of mutated residues are shown as red spheres. The L394P substitution is localized at beta-sheet region far away from the active site. The incorporation of prolyl residue in the mutant probably causes destabilization of secondary structure leading to decreased structural stability of native protein structure.

**Supplementary Figure S4** **Localization of AGAL transiently expressed in HEK 293 cells**

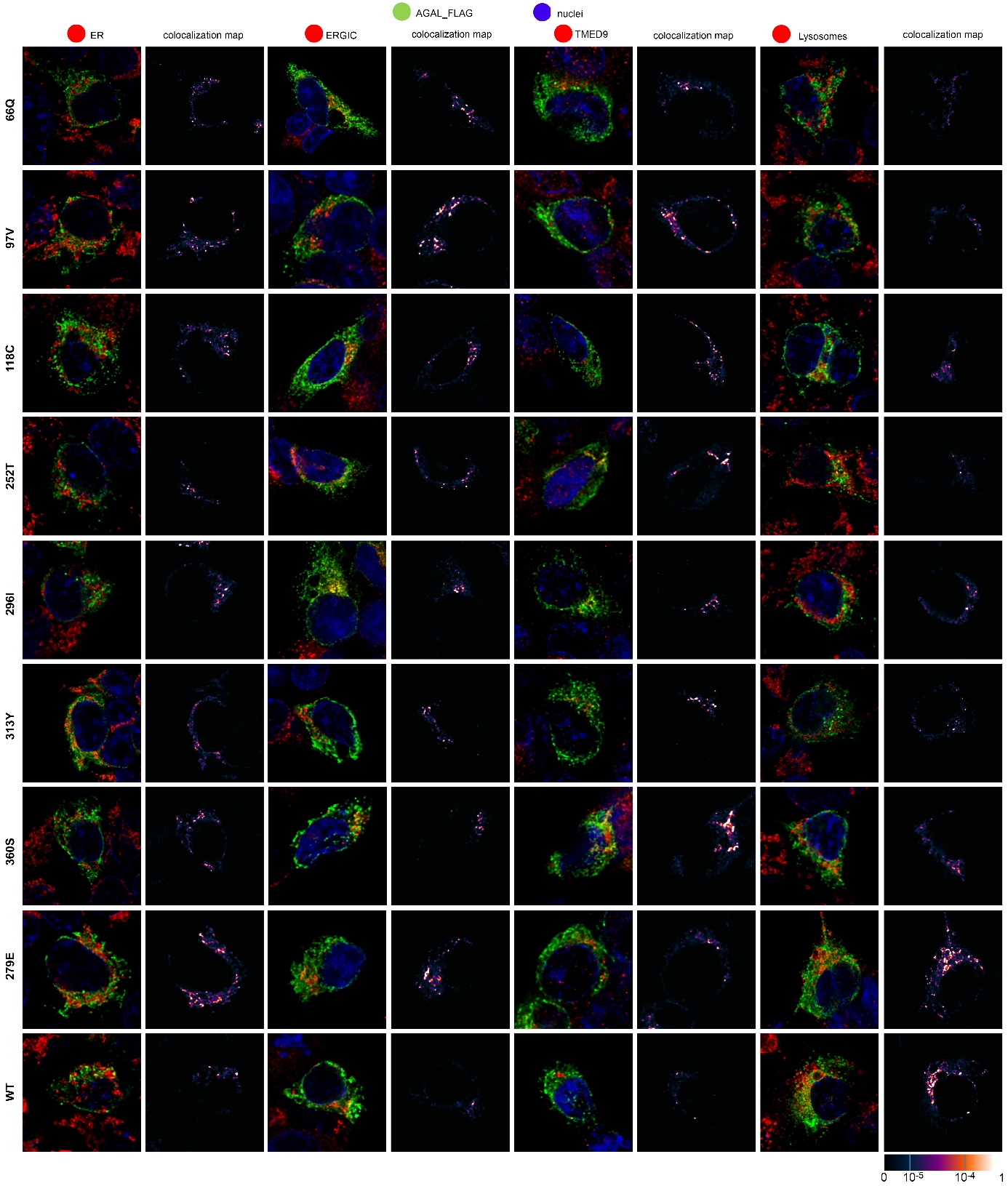

To assess whether and how identified AGAL variants affect intracellular localization, we detected AGAL_FLAG (green) and co-localized the resulting immunofluorescent signal with markers for endoplasmic reticulum (PDI, red), endoplasmic reticulum Golgi intermediate compartment (ERGIC53; red), the secretory cargo receptor TMED9 (red) and lysosomes (LAMP2; red); nuclei are in blue. The degree of AGAL colocalization with selected markers is demonstrated by the fluorescent signal overlap coefficient values that ranges from 0-1. The resulting overlap coefficient values are presented as the pseudo color whose scale is shown in the corresponding lookup tables (LUT).

**Supplementary methods**

**Patients**

The probands were identified from a nationwide Fabry screening program of unselected end-stage kidney disease patients in the Czech Republic. After obtaining informed consent, peripheral blood was drawn and transferred to commercially available filtration paper (CentoCard, CentoGene AG). Samples were dried at room temperature and stored in a plastic sleeve at room temperature for no more than one week and were then sent for analysis. All samples were analyzed at the CentoGene AG laboratory (Rostock, Germany). The algorithm for FD screening was dependent on gender. In males, the AGAL enzymatic activity and the concentration of globotriaosylsphingosine (lyso-Gb3Cer) were determined using fluorimetry and liquid chromatography-mass spectrometry, respectively. If the AGAL activity was < 15.3 µmol/L/h and/or the lysoGb3 concentration >1.8 ng/ml, the entire coding region and the highly conserved exon-intron splice junctions of *GLA* (NM_000169.2) were PCR amplified and Illumina sequenced. In females, given the high rate of expected false-negative results of enzymatic assays, *GLA* sequencing was the first method used. If a variant genotype was found, then the measurement of lyso-Gb3 was done. The result of FD screening was sent to the referring physician. Patients with positive screening results were referred from respective hemodialysis centers to a specialized Fabry Disease center in the General Faculty Hospital, Prague, for further clinical and laboratory investigations and management, including targeted family screening. Patients were evaluated for pre-specified clinical manifestations of Fabry disease including: significant reduction of eGFR (<60 ml/min/1.73m2 bx CKD-EPI formula) or end-stage renal disease requiring renal replacement therapy, presence of microalbuminuria or proteinuria, left ventricular hypertrophy (LVH) detectešd either by echocardiography or cardiac MRI, gastrointestinal signs and symptoms, polyneuropathy, ocular findings (cornea verticillata, Fabry cataract, vessel tortuosities), history of stroke or transitory ischemic attack, and angiokeratomas.

**Biochemical analyses**

AGAL activity in plasma and leukocytes was measured by a fluorometric method using 4-methylumbelliferyl-α-D-galactopyranoside as the substrate (final concentration 2.5 mM ) in the presence of N-acetylgalactosamine (final concentration 0.1 M) as an inhibitor of α-D-galactosidase B [1]. Serum creatinine and other biochemical parameters were measured using routine clinical methods.

**LC-MS analysis of lyso-Gb3Cer, Gb3Cer and met-Gb3Cer**

Plasma lyso-Gb3Cer was extracted using the modified method of Aerts et al [2]. The 50μl of well vortex mixed and sonicated (10s 40J sonication using needle sonicator) plasma was transferred to polypropylene Eppendorf tube and combined with 10μl of chloroform:methanol (2:1, v/v) containing 25ng of D7-d18:1-lyso-Gb3Cer internal standard and 25μl MilliQ water. The mixture was extracted by addition of 450μl of chloroform:methanol (1:2, v/v) followed by 15 min of vortex mixing (1400 RPM) and 10 min of centrifugation at 14 000 x g. The supernatant was transferred to another Eppendorf tube and extracted by the addition of 150μl of chloroform and 225μl of MilliQ water. The mixture was vortex mixed for 10 min (1400 RPM) and then centrifuged for 5 min at 14 000 x g. The upper phase was transferred to new Eppendorf tube and the lower phase was re-extracted with 600μl methanol:water (1:1, v/v) via 10 min vortex mixing (1400 RPM) and 5 min of centrifugation at 14 000 x g. Both upper phases were combined and dried under nitrogen stream. Dried primary extract was then reconstituted in 500μl of MilliQ water and sonicated for 5 min in a sonic bath. Finally, the reconstituted extract was twice extracted using 500μl of water saturated 1-buthanol by 10 min of vortex mixing (1400 RPM) and 5 min centrifugation at 14 000 x g. Both buthanol extracts were combined, transferred to glass vial, and dried under the stream of nitrogen. The final extract was stored at -20oC prior the analysis.

Urinary Gb3Cer and met-Gb3Cer was extracted using the modified method of Abaoui et al [3]. 90μl wasl vortex mixed and sonicated (10 s 40 J sonication using a needle sonicator) urine was transferred to a polypropylene Eppendorf tube and combined with 10μl of chloroform:methanol (2:1, v/v) containing 25ng of (D3-C18:0, d18:1) Gb3Cer internal standard. The mixture was dried under nitrogen stream. 270 μl of methanol and 900 μl of methyl-tert-buthyl ether were added and vortex mixed for 1 min at 1400 RPM, followed by 5 min sonication in a sonic bath, and another 1 min vortex mixing (1400 RPM) and then left standing at laboratory temperature for 1 hour. The 270μl of MilliQ water was added and followed by 1 min of vortex mixing (1400 RPM), a 5 min pause, another 1 min vortex mixing (1400 RPM) and 10 min left standing at laboratory temperature. The mixture was then centrifuged for 5 min at 14 000 x g and after that the samples were placed in a -80^o^C freezer for 40 min.. The lower water phase froze whereas the upper organic phase remained liquid. The organic phase was moved to a glass vial and dried under nitrogen stream. The final extract was stored at -20^o^C prior the analysis.

The lyso-Gb3Cer, Gb3Cer and met-Gb3Cer extracts were reconstituted in 200μl of methanol with 5mM ammonium acetate and filtered through the hydrophobic 0,45μm PTFE syringe filters. The samples were analyzed on a AB/MDS SCIEX API4000 triple quadrupole tandem mass spectrometer with electrospray ionization coupled to the Agillent 1290 Infinity UPLC system equipped with an autosampler. Prior to ionization, the lipids undergo the normal phase LC separation on a Microsolv - Cogent Silica-C 4μm 100A (75mm x 2,1mm) column using methanol with 5mM ammonium acetate mobile phase in isocratic regime at 150 μl/min flow rate. The mass spectrometry was performed in positive ion mode and SRM where [M+H^+^]^+^ precursor ions were generated and fragmented via CID using nitrogen as the collision gas. The product ions with m/z of 282 (lyso-Gb3Cer), 289 (D7-lyso-Gb3Cer IST) and corresponding ceramide for Gb3Cer and met-Gb3Cer were used for quantitative analysis. The quantity was calculated using external calibrator as previously used [4]. The calibrator consists of 25ng of D7-lyso-Gb3Cer or (D3-C18:0, d18:1) Gb3Cer internal standards combined with 50ng of lyso-Gb3Cer or Gb3Cer according to quantified lipid.

The mass spectrometer electrospray and ion optics setting were as follows for analysis of lyso-Gb3Cer - CAD 7psi, CUR 20psi, GS1 30psi, GS2 50psi, IS 5500V, TEM 200oC, ihe ON, DP 120V, EP 10V, CE 52V, CXP 12V for Gb3Cer and met-Gb3Cer the setting were – CAD 7psi, CUR 20psi, GS1 30psi, GS2 50psi, IS 5500V, TEM 200oC, ihe ON, DP 135V, EP 11V, CE 43V, CXP 16V.

The product and precursor ion transition pairs for Gb3Cer and met-Gb3Cer were:

Precursor (Da)	product (Da)	Dwell(msec)	ID

1024,6	520,2	200	C16-Gb3Cer

1052,7	548,3	200	C18-Gb3Cer

1080,8	576,4	200	C20-Gb3Cer

1106,8	602,4	200	C22:1-Gb3Cer

1108,7	604,3	200	C22-Gb3Cer

1134,8	630,4	200	C24:1-Gb3Cer

1136,8	632,4	200	C24-Gb3Cer

1038,8	534,4	200	C16Met-Gb3Cer

1066,8	562,4	200	C18Met-Gb3Cer

1094,8	590,4	200	C20Met-Gb3Cer

1120,9	616,5	200	C22:1Met-Gb3Cer

1122,9	618,5	200	C22Met-Gb3Cer

1148,9	644,5	200	C24:1Met-Gb3Cer

1150,9	646,5	200	C24Met-Gb3Cer

**Degradation of globotriaosylceramide in cultured skin fibroblasts**

Skin fibroblasts were grown in Dulbecco’s modified Eagle’s medium (DMEM) supplemented with 10% fetal calf serum (FCS), glutamine and pyruvate at 37°C in an incubator with 5% CO2. After achieving 85% confluence, the cells were loaded with mass labeled C23:0,d18:1 globotriaosylceramide (C23Gb3Cer) to a final concentration of 3µg/mL media. Conduritol B epoxide (Calbiochem-Novabiochem GmbH, Germany), a covalent inhibitor of GlcCer-β-Glucosidase, was added to a final concentration of 0.5 mM to block the downstream metabolic conversion of glucosylceramide to ceramide. After 4 days, the medium was removed, cells were harvested using trypsin, washed in PBS and sonicated using a Branson cup-horn sonicator for 30 seconds at 20% output. Sphingolipids were extracted from the cell homogenate (10µg of protein aliquote) using 1ml of chloroform:methanol (2:1; v/v) and 50 ng of the internal standard C17:0 globotriaosylceramide. After extraction, the samples were dissolved in 1ml of methanol:water (4:1; v/v) with 5mM ammonium acetate and quantified using the flow injection analysis electrospray ionization tandem mass spectrometry (FIA-ESI-MS/MS) on AB/MDS SCIEX API4000 tripple quadrupole tandem mass spectrometer coupled to an UPLC Agilent 1290 Infinity equiped with an autosampler. The samples were injected into the methanol:water (4:1; v/v) mobile phase with a flow rate of 50ul/min. Other conditions were as previously described [4, 5].

**Intracellular localization of AGAL in kidney biopsies**

For intracellular localization of AGAL in kidney biopsy we stained always in parallel AGAL and LAMP-2 marker of lysosomes with markers of the ER, ERGIC or Golgi. AGAL was detected with the rabbit monoclonal anti-galactosidase alpha antibody (Abcam, ab168341, Cambridge, UK), diluted 1:500. Lysosomes were detected with goat polyclonal anti-human LAMP-2/CD107b antibody (R and D systems, AF6228, Minneapolis, USA) diluted 1:50. Endoplasmic reticulum was detected with mouse anti-PDI (ADI-SPA-891, Enzo, Farmingale, NY, USA), diluted 1:50. The ERGIC was detected with mouse monoclonal anti-LMAN1 antibody (clone OTI1C5, Invitrogen, Thermo Fisher Scientific, Waltham, MA, USA), diluted 1:100. The Golgi apparatus was detected by mouse anti-Golgi 58K protein /formiminotransferase cyclodeaminase (FTCD) antibody (G2404, SIGMA-Aldrich, Prague, Czech Republic). The incubation of primary antibodies was overnight at 4°C.

Detection of bound primary antibodies was achieved using donkey anti-rabbit IgG Alexa Fluor 555, donkey anti-goat IgG Alexa Fluor 488 or donkey anti-mouse IgG Alexa Fluor 647 secondary antibodies (Thermo Fischer Scientific) diluted 1 : 500 in 5% BSA in PBS and incubated 1 hour at 37°C.

Colocalization of AGAL, lysosome and TMED9: For detection of AGAL and lysosome, primary antibodies were diluted as described above and incubated for 1h at 37°C, followed by incubation with secondary antibodies (as above) for 1h at 37°C. After a quick wash, rabbit polyclonal anti-TMED9 antibody (21620-1-AP, Proteintech, Manchester, UK) directly labeled with Dylight 488 Fast Conjugation Kit (Abcam, Cambridge, UK) was added. TMED9-AF488 labeled antibody was diluted 1 : 50 and incubated overnight at 4°C.

**Intracellular localization of AGAL in primary skin fibroblasts**

For intracellular localization of AGAL in primary skin fibroblasts, we stained in parallel AGAL and individual compartments- ER, ERGIC, Golgi, TMED9 and lysosome, respectively.

AGAL was detected as described above with antibody diluted 1:300. For detection of endoplasmic reticulum we used mouse anti-PDI (specified above) diluted 1:200, ERGIC was detected with mouse monoclonal anti-LMAN1 (specified above) diluted 1:200. Lysosomes were detected with mouse monoclonal anti-H4B4 antibody (Ab25631, Abcam) diluted 1:200. The incubation of primary antibodies was overnight at 4°C.

Detection of bound primary antibodies were achieved using donkey anti-mouse IgG Alexa Fluor 488 donkey anti-rabbit IgG Alexa Fluor 555 secondary antibodies (Thermo Fischer Scientific) diluted 1: 1000 and incubated 1 hour at 37°C. Slides were mounted in ProLong gold antifade mountant with DAPI (Thermo Fischer Scientific ) and analyzed by confocal microscopy.

Colocalization of AGAL and TMED9: For parallel staining of AGAL and TMED9, AGAL was first detected by rabbit monoclonal anti-galactosidase alpha antibody (Abcam, ab168341, Cambridge, UK) diluted 1:200, incubated overnight at 4°C, followed by 1 hour incubation with secondary antibody anti-Rabbit IgG conjugated AF 555 (Thermo Fisher Scientific, Waltham, MA, USA) at 37°C diluted 1:1000. After a quick wash, the cells were incubated overnight at 4°C with rabbit anti-TMED9 antibody directly labeled with Alexa Fluor 488, diluted 1 : 20.

Slides were mounted in the fluorescence mounting medium ProLong gold antifade mountant with DAPI (Molecular Probes, Invitrogen) and analyzed by confocal microscopy as described above.

**Intracellular localization of AGAL in HEK293 cells stably or transiently transfected with *GLA_FLAG cDNA***

Wild-type *GLA* cDNA was reverse transcribed from human leucocyte total RNA, polymerase chain reaction (PCR) amplified, cloned into pCR- XL-TOPO vector and introduced into the One Shot® TOP10 Chemically Competent *E. coli* (Invitrogen, Paisley, UK). A single 3-terminal flag-tag sequence was appended to the originally synthesized wild type *GLA* cDNA using PCR with specific oligonucleotide primers containing the BamHI and Xho I restriction sites for subsequent subcloning of the *GLA_FLAG* construct to pCR3.1 vector (Invitrogen, Paisley, UK). The resulting product *GLA_FLAG/*pCR3.1 was introduced into the Escherichia coli TOP 10´F strain and clones with correct *GLA* sequence were selected by Sanger sequencing. Mutant constructs (mut_*GLA_FLAG*/pCR3.1) were prepared by site-directed mutagenesis (Site-Directed Mutagenesis Kits,QuikChange II, Agilent). and selected by sequencing.

For stable AGAL_FLAG expression, HEK 293 cells were maintained in Dulbecco's Modified Eagle Medium : Nutrient Mixture F-12 (DMEM/F-12) high glucose medium supplemented with 10% (vol/vol) fetal calf serum (Gibco), 100 U/ml penicillin G/streptomycin (Sigma, Prague, Czech Republic). HEK293 cells were transfected with 2.5 µg of plasmid DNA using Lipofectamine 3000^TM^ (Invitrogen, Paisley, UK). Three days post transfection, cells were trypsinized, diluted and cultured in selective medium containing 0.5 mg/ml G418 (Invitrogen-Gibco, Paisley, UK). For each selected clone, the presence and correct sequence of *GLA_FLAG* was confirmed by Sanger sequencing; the amount of *GLA*_*FLAG* transcript was determined by real-time PCR, and expression levels of AGAL_FLAG tagged proteins were assesed by Western blot with mouse monoclonal anti-FLAG antibody (F1804, SIGMA-Aldrich, Prague, Czech Republic). Clones demonstrating similar *GLA_FLAG* transcript and AGAL_FLAG protein amounts were selected for further analyses.

For transient AGAL_FLAG expression, transfections were carried out using Lipofectamine 3000TM (Invitrogen, Paisley, UK) with either 0.5 µg or 2,5µg DNA for 1.5x10^5^ or 8x10^5^ cells, respectively. Expression of AGAL_FLAG proteins was assessed 24 hours post transfection by Western blot analysis with mouse monoclonal anti-FLAG antibody (F1804, SIGMA-Aldrich, Prague, Czech Republic). Endogenously expressed AGAL and tubulin were detected with rabbit monoclonal anti-galactosidase alpha antibody (used for staining of AGAL in kidney), mouse monoclonal anti-acetylated tubulin antibody (T7451, SIGMA-Aldrich, Prague, Czech Republic), respectively.

For intracellular localization in stably or transiently transfected HEK293 cells, we stained in parallel AGAL_FLAG and lysosomes with markers of the individual compartments ER, ERGIC, Golgi and TMED9, respectively. AGAL_FLAG was detected with mouse monoclonal ANTI-FLAG® M2 antibody (F1804, Sigma Aldrich) diluted 1:100. Lysosomes were detected with rabbit polyclonal anti-LAMP-2 (customly prepared) diluted 1:4000. The endoplasmic reticulum was detected with rabbit polyclonal anti-SEC61A1 antibody (ab183046, Abcam) diluted 1:500, ERGIC was detected with rabbit polyclonal anti-LMAN1 (ProteinTech) diluted 1:300. The Golgi apparatus was detected by rabbit monoclonal anti-GM130 (C-terminal, G7295, SIGMA-Aldrich, Prague, Czech Republic) diluted 1:300. TMED9 protein was detected with rabbit polyclonal TMED9 antibody (21620-1-AP, Proteintech) diluted 1:100. The primary antibodies were incubated overnight at 4°C.

Detection of bound primary antibodies was achieved using donkey anti-mouse IgG Alexa Fluor 488 donkey anti-rabbit IgG Alexa Fluor 555 secondary antibodies (Thermo Fischer Scientific) diluted 1: 1000 and incubated for 1 hour at 37°C. Slides were mounted in ProLong gold antifade mountant with DAPI (Thermo Fischer Scientific) and analyzed by confocal microscopy as described above.

**AGAL secretion**

Secreted AGAL was detected in media of transiently transfected HEK293 cells (see above). Media were processed as described before (Živná et al., Kidney Int. 2020 Dec; 98(6): 1589–1604, Supplemental Material). Here, concentrated media were resolved on 12% separating gel in SE640apparatus (Hoefer, San Francisco, CA, USA). Proteins were transferred Immobilon-E PVDF membrane (Merck Millipore, Tullagreen, Ireland) in semi-dry blotting apparatus Trans-Blot® SD Cell (Bio-Rad, Prague, Czech Republic) at 0.6 mA/cm2 for 60 minutes. Membrane was blocked in Tris-buffered saline (TBS) and 5% non-fat milk for 2 h at room tempereature (RT) followed by 3x10 min washes in TBS. AGAL was detected by monoclonal Anti-Galactosidase alpha antibody (# ab168341, Abcam, Cambridge, UK) diluted 1:2 000 in TBS with 0.1% Tween 20 (TBST) and 2% non-fat milk for 1 h/RT followed by 4x10 min washes in TBST. Primary antibody was detected by HRP-conjugated Goat anti-Rabbit IgG (H+L) secondary antibody (# 31460, Thermo Fisher Scientific, Prague, Czech Republic) diluted 1:10 000 in TBST + 2% non-fat milk for 1 h/RT followed by 3x10 min washes in TBST and 1x5 min in TBS. Membranes were then incubated in Clarity Max™ Western ECL Substrate (Bio-Rad, Prague, Czech Republic) according to manufacturer’s instructions. Chemiluminiscent signal was recorded by ChemiDoc™ MP Imaging System (Bio-Rad, Prague, Czech Republic).

**RNA sequencing of HEK293 cells stably expressing AGAL_FLAG**

Total RNA was isolated from stably transfected cells using the RNA Minikit (Qiagen). RNA concentrations were determined spectrophotometrically at A260 nm by NanoDrop (NanoDrop Technologies, Wilmington, DE), and RNA quality was verified using an Agilent 2100 bioanalyser - RNA Lab-On-a-Chip (Agilent Technologies, Santa Clara, USA). Aliquots of isolated RNA were stored at –80°C until analysis. A stranded mRNA-Seq library was prepared with KAPA mRNA HyperPrep Kit for Illumina® Platforms (Roche). Paired-ends reads of 2x100 base pairs were sequenced on Illumina NoveSeq6000 at the National Center fo Medical Genomics in Prague according to manufacturer protocol. The resulting FASTQ files was subjected to QC control and trimmed using atropos (v.1.128)[6]. Gene-level abundance were estimated using Salmon (v.1.3) [7] with the Ensembl gene definition v.75. Normalization and differential expression analyses were performed with the DESeq2 R package [8]. The accession number for the RNA-Seq data reported in this paper is GEO: XXXXXXXXXX

**Proteomic analysis of HEK293 cells stably expressing AGAL_FLAG**

Cell pellets were homogenized and lysed by boiling at 95°C for 10 min in 100mM TEAB (Triethylammonium bicarbonate) containing 2% SDC (sodium deoxycholate), 40mM chloroacetamide, 10mM TCEP (Tris(2-carboxyethyl)phosphine) and further sonicated (Bandelin Sonoplus Mini 20, MS 1.5). Protein concentration was determined using the BCA protein assay kit (Thermo), and 30 µg of protein per sample was used for MS sample preparation.

Samples were further processed using SP3 beads according to Hughes et al. [9]. Briefly, 5 µl of SP3 beads was added to 30 µg of protein in lysis buffer and filled to 50 µl with 100mM TEAB. Protein binding was induced by addition of ethanol to 60 % (vol./vol.) final concentration. Samples were mixed and incubated for 5 min at room temperature. After binding, the tubes were placed in a magnetic rack, and the unbound superntatant was discarded. Beads were subsequently washed twice with 180 µl of 80% ethanol. After washing, samples were digested with trypsin (trypsin/protein ratio 1/30) and reconstituted in 100mM TEAB at 37°C overnight. After digestion, samples were acidified with TFA to 1% final concentration, and peptides were desalted using in-house made stage tips packed with C18 disks (Empore), according to Rappsilber et al. [10]

nLC-MS 2 Analysis

Nano reversed phase columns (EASY-Spray column, 50 cm x 75 µm ID, PepMap C18, 2 µm particles, 100 Å pore size) were used for LC/MS analysis. Mobile phase buffer A was composed of water and 0.1% formic acid . Mobile phase B was composed of acetonitrile and 0.1% formic acid. Samples were loaded onto the trap column (C18 PepMap100, 5 μm particle size, 300 μm x 5 mm, Thermo Scientific) for 4 min at 18 μl/min loading buffer was composed of water, 2% acetonitrile and 0.1% trifluoroacetic acid . Peptides were eluted with a mobile phase B gradient from 4% to 35% B in 120 min. Eluting peptide cations were converted to gas-phase ions by electrospray ionization and analyzed on a Thermo Orbitrap Fusion (Q-OT- qIT, Thermo Scientific). Survey scans of peptide precursors from 350 to 1400 m/z were performed in Orbitrap at 120K resolution (at 200 m/z) with a 5 × 10^5^ ion count target. Tandem MS was performed by isolation at 1,5 Th with the quadrupole, HCD fragmentation with normalized collision energy of 30, and rapid scan MS analysis in the ion trap. The MS2 ion count target was set to 10^4^, and the maximum injection time was 35 ms. Only those precursors with charge state 2–6 were sampled for MS2. The dynamic exclusion duration was set to 45 s with a 10 ppm tolerance around the selected precursor and its isotopes. Monoisotopic precursor selection was turned on. The instrument was run in top speed mode with 2 s cycles.

Data analysis

All data were analyzed and quantified with the MaxQuant software (version 1.6.3.4) [11]. The false discovery rate (FDR) was set to 1% for both proteins and peptides and we specified a minimum peptide length of seven amino acids. The Andromeda search engine was used for the MS/MS spectra search against the human database (downloaded from Uniprot on August 2020, containing 20 365 entries). Enzyme specificity was set as C-terminal to Arg and Lys, also allowing cleavage at proline bonds and a maximum of two missed cleavages. Dithiomethylation of cysteine was selected as a fixed modification and N- terminal protein acetylation and methionine oxidation as variable modifications. The “match between runs” feature of MaxQuant was used to transfer identifications to other LC-MS/MS runs based on their masses and retention time (maximum deviation 0.7 min) and this was also used in quantification experiments. Quantifications were performed with the label-free algorithm in MaxQuant [12]. Data analysis was performed using Perseus 1.6.1.3 software [13].

Pathway analysis of the proteomic and RNASeq data was performed with GSEA (v4.1.0). Differentially expressed proteins were categorized on two different levels, canonical pathways (CP) and gene ontology (GO) terms (C2 and C5 gene set collections in MSigDB v7.2). The terms with FDR q-value <0.05 were visualized using Enrichment map v3.3.3 plugin in Cytoscape v3.8.2.

UPR branch activation analysis is presented as boxplots. Each box of the boxplot consists of proteins comprising an individual UPR branch as defined [14]. The expression of each protein in the group is plotted as mean of the scaled expression profiles obtained from three replicates as described [15].

**qRT-PCR analysis of UPR markers in HEK293 cells transiently transfected with *GLA_FLAG cDNA***

RNA was prepared from transiently transfected HEK293 cells using RNeasy Mini Kit (Qiagen) or TRIzol reagent (Invitrogen) according to the manufacturer´s protocol. Samples of 15 ng total RNA were reverse trancribed and amplified using SOLIScript 1-step Multiplex Probe Kit (Solis Biodyne) according to the manufacturer´s instructions. PCR master mix contained 300 nM of each primer and TaqMan probe (Generi Biotech) in a total reaction volume of 20 µl. The qPCR reactions were run on a CFX96 instrument (Bio-Rad). Relative expression levels were calculated by 2^–∆∆Ct^ method using *PUM1*, *CASC3*, *UBC* and *TBP* genes as a reference. Primer and probe sequences used are listed in the following table.

| Probe/primer name | Sequence |
| --- | --- |
| HSP90B1_probe | /56-FAM/CGT TCC CCG /ZEN/TCC TAG AGT GTT TCC /3IABkFQ/ |
| PDIA4_probe | /56-FAM/CAT GGT GTG /ZEN/GAC ATT GCA AGC AGT /3IABkFQ/ |
| HYOU1_probe | /56-FAM/TGC ATC CCA /ZEN/GCT TCC TTA GTC TTC AC/3IABkFQ/ |
| CRELD2_probe | /56-FAM/TGA AAG TGT /ZEN/GCT GCT CTC CAG GAA C/3IABkFQ/ |
| MANF_probe | /56-FAM/CAC AAC CGA /ZEN/TTC TCT TTG CCT CTT GC/3IABkFQ/ |
| HSPA5_probe | /56-FAM/CCC AAC GCC /ZEN/AAG CAA CCA AAG AC/3IABkFQ/ |
| SEC61A1_probe | /56-FAM/ACT GTC AAC /ZEN/ACT GGC CGA GGA AT/3IABkFQ/ |
| TMED9_probe | /56-FAM/AGT GAA AGT /ZEN/GAA CCT GCC CTC GG/3IABkFQ/ |
| XBP1s_probe | /56-FAM/CTG AGT CCG /ZEN/CAG CAG GTG CA/3IABkFQ/ |
| XBP1u_probe | /5HEX/AGC ACT CAG /ZEN/ACT ACG TGC ACC TCT /3IABkFQ/ |
| SAA1_1_probe | /5HEX/AGG CTG CCA /ZEN/ATG AAT GGG GCA /3IABkFQ/ |
| PUM1_probe | /5Cy5/AGC AGC AAC /TAO/TGT GGG ACT TTT TGA C/3IAbRQSp/ |
| TBP_probe | /5Cy5/TGG GAT TAT /TAO/ATT CGG CGT TTC GGG C/3IAbRQSp/ |
| CASC3_probe | /5Cy5/TGC CCC GGA /TAO/CTC CAA TTC TGT T/3IAbRQSp/ |
| POLR2A_probe | /5Cy5/TCA CAG ACA /TAO/TTC GCT TCA GTT CAT CCG /3IAbRQSp/ |
| UBC_probe | /5Cy5/TCG TGA AGA CTC TGA CTG GTA AGA CC /BHQ2/ |
| HSP90B1_FOR | AGC ACA TCT GGG AGT CTG A |
| PDIA4_FOR | CTG ACA AAG ACA CAG TGC TG |
| HYOU1_FOR | CGC CGG AAA GAT ATT AAC ACC |
| CRELD2_FOR | CTG ACT TAT TCG AGT GGT TTT GTG |
| MANF_FOR | GGG AAG ATT TTA CCA GGA CCT C |
| HSPA5_FOR | GAA ACC GCT GAG GCT TAT TTG |
| SEC61A1_FOR | TCG TAT GGA AGG CAT TCA GC |
| TMED9_FOR | TGT TTG TGG AGG TGA AGG AC |
| PUM1_FOR | TTA TTC AGG CAC GCA GGT AC |
| TBP_FOR | GAG AGT TCT GGG ATT GTA CCG |
| CASC3_FOR | CCG GTG GAT TCT AGT ACA AGT G |
| POLR2A_FOR | CCA TCA AGA GAG TCC AGT TCG |
| HSP90B1_REV | CCA ATT CAA GGT AAT CAG ATG CTT C |
| PDIA4_REV | TGA GGT TGC ATC GAT CTT GG |
| HYOU1_REV | GGG TAC GGT CAA ATC CTA CTC |
| CRELD2_REV | TGT CTG CTC CCA TCT CCG |
| MANF_REV | CAT CTG TGG CCC CGA TAT AG |
| HSPA5_REV | CTG CCG TAG GCT CGT TGA T |
| SEC61A1_REV | GGG AAG ATT CTG GCG GTA G |
| TMED9_REV | AGC AAA GAG GGA GAA CTT GG |
| PUM1_REV | CGC ATT AGG TCT TTG GAA CAG |
| TBP_REV | ATC CTC ATG ATT ACC GCA GC |
| CASC3_REV | GTC CTG CTC CCA TGT GTA TAT G |
| POLR2A_REV | CCA GTC CGC TCA ATC ACC |
| UBC_FOR | GAT CGC TGT GAT CGT CAC TTG |
| UBC_REV | GTT TTC CAG CAA AGA TCA GCC T |
| XBP1_FOR | CAT GGC CTT GTA GTT GAG AAC |
| XBP1_REV | AGA ATG CCC AAC AGG ATA TCA |

**Supplementary Table S3.** AGAL variants of unknown significance and mutations causing FD type 1 and 2 characterized in this study

| AGAL  variant | Prague database | AGAL-leucocytes  [nmol.mg-1.h-1]  Controls; (n=477)  mean±SD  59.7±14.6  Range  24.8-103 | plasma AGAL  [nmol.ml-1.h-1]  Controls; (n=322)  mean±SD  6.1±2.8  Range  2.4-19.4 | organs affected | pathologic changes | Pathogenicity * | population frequency ** | references |
| --- | --- | --- | --- | --- | --- | --- | --- | --- |
| **Variants of unknown significance** | | | | | | | | |
| p.L394P | Yes  n=17 | M 8.5 ± 1.0 (n=8)  F 53.1 ± 18.9 (n=9) | M 0.9 ± 0.3 (n=8)  F 4.6 ± 3.6 (n=9) | kidney | interstitial fibrosis;  podocytopathy with lipoid nephrosis | n.a. | 0 | this report |
| p.A143T | Yes  n= 12 | M 17.2 ± 5.4 (n=5) F 40.2 ± 12.5 (n=7) | M 1 ± 0.3 (n=5) F 2.5 ± 1.1 (n=6) | kidney  OR  brain | interstitial fibrosis;  podocytopathy with lipoid nephrosis | likely benign | 5.1 x 10^-4^ | The genetic cause of the initial report of FD [1]. Low to normal AGAL activity, no increase in plasma Gb3Cer and lysoGb3Cer; lysosomal storage vacuoles are usually absent in affected organs; heterogeneous clinical presentation with multiple clinically unaffected carriers [2-8] |
| p.E66Q | no | n.a. | n.a. | n.a. | n.a. | benign | 1.1 x 10^-4^ | Low to normal AGAL activity, no increase in plasma Gb3Cer and lysoGb3Cer; predominantly or solely affect kidney, zebra bodies absent in affected kidney [9-16] |
| p.A97V | no | n.a. | n.a. | n.a. | n.a. | Later onset | 0 | Low to normal AGAL activity, no increase in plasma Gb3Cer and lysoGb3Cer; heterogeneous clinical presentation with multiple clinically unaffected carriers [17, 18] |
| p.R118C | Yes  n=3 | F 44.3 ± 14.5 (n=3) | F 2.0 ± 1.25 (n=3) | Brain  AND  eye | n.a. | benign | 2.3 x 10^-4^ | normal Lyso-Gb3Cer, Fabry-specific renal pathology absent [19, 20] |
| p.R252T | Yes  n=2 | M 80.6 F 72.1 | M 14.2 F 5.98 | kidney | n.a. | benign | 1.9 x 10^-5^ | normal Lyso-Gb3Cer [8], |
| p.M296I | no | n.a. | n.a. | n.a. | n.a. | Later onset | 0 | Residual activity AGAL activity, normal Lyso-Gb3Cer [21], progressive renal disease with absence of Fabry-specific renal pathology [17, 22], originally identified in case with left ventricular hypertrophy and presence of typical lysosomal storage vacuoles in myocardial cells [23] |
| p.D313Y | Yes  n=13 | M 26.4 ± 5.2 (n=3) F 35.6 ± 10.7 (n=8) | M 1.2 ± 0.5 (n=5) F 2.2 ± 0.8 (n=8) | brain  OR kidney | n.a. | benign | 3 x 10^-3^ | Originally identified in classically affected males. Low to normal AGAL activity, plasma Gb3Cer and lysoGb3Cer normal in majority of cases, with absence of Fabry-specific renal pathology [24], recent studies suggest late onset neuropathy involvement [2, 24, 25]. Pathogenicity still debated [26]. |
| **Mutations causing classic FD type 1 and 2** | | | | | | | | |
| p.G360S | Yes  n=6 | M 2.1 ± 0.2 (n=2) F 17.2 ± 5.3 (n=3) | M 1.0 ± 0.21 (n=2) F 3.2 ± 0.37 (n=3) | kidney, heart, brain, eye | multilamellar lysosomal storage vacuoles | classic | 0 | [27] |
| p.Q279E | no | n.a. | n.a. | n.a. | n.a. | Later onset | 0 | Severely reduced AGAL-A activity due to reduced protein stability [28] |
| p.R112H | Yes  n=2 | M 0.79 F 16.5 | M 0.83 F 1.86 | kidney, heart | n.a. | pathogenic, various onset | 1.1 x 10^-5^ | Residual activity AGAL activity and heterogeneous phenotype [6, 18, 29, 30], no increase in plasma Gb3Cer and lysoGb3Cer [31]; zebra bodies present in affected kidney [30, 32, 33] |
| p.A135V | Yes  n=2 | M 2.8  F 55.6 | M 0.68 F nd | kidney, heart, brain | multilamellar lysosomal storage vacuoles | classic | 0 | [27] |
| p.R301Q | Yes  n=6 | M 2.4 ± 0.5 (n=2) F 22.9 | M 0.86 ± 0.12 (n=2) F 1.95 | kidney, heart | multilamellar lysosomal storage vacuoles | pathogenic, later onset | 0 | Severely reduced AGAL-A activity, increase in plasma Gb3Cer and lysoGb3Cer; lysosomal storage vacuoles are present in affected organs; cardiac and renal manifestations [28, 34-37] |
| p.R342Q | Yes  n=22 | M 1.7 ± 0.8 (n=8) F 29.1 ± 14.4 (n=14) | M 0.2 ± 0.04 (n=6) F 2.6 ± 1.9 (n=12) | kidney, heart, brain | multilamellar lysosomal storage vacuoles | classic | 0 | [27] |

M – males; F-females; * International Fabry Disease Genotype-Phenotype Database (dbFGP) available at <http://www.dbfgp.org/dbFgp/fabry/>; ** GnomAD, Genome aggregation database, version v2.1.1 available at https://gnomad.broadinstitute.org/; n.a. not availbale
